# A humoral immune response to parasitoid wasps in *Drosophila* is regulated by JAK/STAT, NF-κB and GATA

**DOI:** 10.1101/2024.06.12.598701

**Authors:** Shuyu Olivia Zhou, Jonathan P. Day, Bart Deplancke, Alexandre B. Leitão, Francis M. Jiggins

**Author notes:** **Corresponding Authors:** Shuyu Olivia Zhou, Alexandre B. Leitão, Francis M. Jiggins, Address for correspondence: Francis M. Jiggins, Department of Genetics, University of Cambridge, Downing Pl, Cambridge, CB2 3EH, UK, Phone number for correspondence.

## Abstract

The two arms of innate immunity consist of the cell-mediated cellular defenses and the systemic humoral immune responses. *Drosophila* humoral immune defenses in the context of antimicrobial immunity, particularly the regulation and activation of antimicrobial peptide secretion from the fat body, have been studied extensively. How *Drosophila* regulates humoral immunity against another major natural enemy, the parasitoid wasp, is less well-characterized. In this study, we focused on a gene crucial in anti-parasitoid immunity, *lectin-24A*, which is specifically induced following parasitization. We found that a fluorescent reporter driven by the region upstream of *lectin-24A* showed localized posterior expression in the larval fat body, the *Drosophila* tissue mediating humoral immunity. Furthermore, with RNA sequencing of the anterior and posterior fat body sections, we found that components of JAK/STAT, GATA, and Toll pathways were regulated differentially in the anterior-posterior axis of the fat body and/or by infection. Predicted binding motifs for transcription factors in all three of these pathways were identified in the 444bp upstream region of the *lectin-24A* gene, where scrambling these motifs leads to reduced basal or induced expression of the fluorescent reporter. Investigating each of these pathways, we found that JAK/STAT, the GATA factor Pannier, and the NF-κB factor dorsal all modulate the expression of *lectin-24A*. The binding motifs associated with these transcription factors were also enriched in the upstream sequences of parasitism-induced genes in the fat body. Taken together, these results indicate that JAK/STAT, Pannier, and NF-κB signaling are involved in the regulation of *lectin-24A* and, more generally, *Drosophila* humoral anti-parasitoid immunity after infection.

## Introduction

Parasitoid wasps are major natural parasites of insects. In *Drosophila melanogaster*, the most common parasitoids attack larvae. Females of species such as *Leptopilina boulardi* insert an egg via their ovipositor inside the *Drosophila* larva, where the parasitoid larva, once emerged, feeds on host tissue during development. Without a successful immune response, the *Drosophila* host is eventually killed at the pupal stage. However, parasitization induces an anti-parasitoid defense in the larva termed melanotic encapsulation that involves both cellular and humoral immune responses [1–3]. *Drosophila* immune cells (hemocytes) are recruited to the parasitoid egg and surround it in a multi-layered cellular capsule [4–6]. This capsule then becomes melanized, killing the parasitoid [7].

Several humoral immune effectors have been identified in the context of humoral anti-parasitoid immunity in *Drosophila*. Humoral immunity is mediated by the *Drosophila* fat body, an insect-specific organ functionally similar to the mammalian liver, the metabolic hub of the organism. The invertebrate counterparts of the mammalian complement factors, called thioester-containing proteins (TEPs) [8], a group of serine proteases [9,10], and the fat body-derived peptide edin [11], are some of the secreted immune effectors shown to be involved in this response. We have previously also shown that a C-type lectin, lectin-24A, secreted by the larval fat body, plays an important role in this immune response [12]. The induction of *lectin-24A* expression following parasitism, controlled by the *cis*-regulatory upstream region of the gene, is associated with higher resistance to parasitoid infections [12]. Lectin-24A localizes to the surface of the parasitoid egg before hemocytes attach to it, suggesting that it may play a role in the recognition of the egg and subsequent recruitment of hemocytes.

*Drosophila* humoral immunity against bacterial and fungal infections has been investigated intensively in the past and is well characterized. Following recognition of pathogens, humoral defenses are induced via the activation of the two hallmark NF-κB immune signaling pathways, Toll and Imd (reviewed in [13] and [14]). These two pathways are known for their roles in regulating the secretion of many immune-related genes by the fat body, most famously antimicrobial peptides (AMPs). Dif and Dorsal are the two NF-κB family transcription factors (TFs) in the Toll pathway, which is mostly responsible for defense against Gram-positive bacteria, fungi and other pathogens. Relish is the transcription factor involved in Imd signaling, which mostly controls the defense against gram-negative bacteria. In some cases, a combination of the two pathways is responsible for the expression of immune effectors in the fat body [15,16]. Other than its role in the induction of AMP expression, the Toll pathway is also involved in regulating the proliferation of hemocytes in *Drosophila* larvae [17,18]. Loss-of-function mutations in the Toll pathway components result in reduced capacity to melanize parasitoids [13,17], whereas overactivation of the Toll pathway leads to a spontaneous melanization phenotype reminiscent of the melanotic encapsulation response against parasitoid wasps [13,17,19].

The Janus kinase/signal transducer and activator of transcription (JAK/STAT) pathway has been shown to be involved in the cellular immune response through the regulation of hemocyte differentiation following parasitoid infection [18]. Downregulation of JAK/STAT signaling in the anterior lobes of the larval lymph gland plays an important role in regulating the differentiation of lamellocytes, a specialist type of hemocyte for the encapsulation response [20,21]. It was also found that the activation of JAK/STAT signaling in somatic muscles by hemocyte-derived cytokines Upd2 and Upd3 releases energy stored in the muscles as glycogen for use in the immune response and is key to the immune response against wasp infection [22,23]. Similar to the Toll pathway, overactivation of JAK/STAT signaling in the whole larva also results in an over-proliferation of hemocytes and spontaneous melanization [18,24,25].

A third group of transcription factors mediating *Drosophila* immunity is the GATA family of transcription factors, a conserved group of TFs with zinc finger domains that recognize variations of the GATA motif ((A/T) GATA (A/G)) (reviewed in [26]). *Drosophila* has a total of five GATA factors: Pannier, Serpent, Grain, dGATAd, and dGATAe. Serpent is essential for the differentiation of fat body and hemocytes [27,28], and Pannier has been found to play an important role in hemocyte differentiation and maturation [29]. Strikingly, many immunity genes expressed by the fat body exhibit Rel-Serpent synergy, where closely linked NF-κB and GATA binding sites are found in the upstream regulatory region [30,31]. While Serpent is the principal GATA factor regulating the antimicrobial response in the *Drosophila* fat body, other GATA factors were reported to regulate immune responses in other *Drosophila* tissues [31].

The anti-parasitoid humoral response is mainly driven by wasp parasite-associated molecular patterns (PAMPs) [32], but the signaling pathways involved in the response remain unknown. Previous studies of the transcriptional response to parasitoid attack in *Drosophila* larvae have identified a series of genes that are differentially expressed in response to wasp oviposition [33–36]. These studies have implicated the three pathways outlined above in the response [33,35]. In particular, Wertheim et al. found that transcription factor binding motifs (TFBMs) for NF-κB, STAT, and GATA significantly associate with genes differentially expressed after parasitism [33]. However, these observations were not followed up experimentally. Previous investigations have also focused on transcriptional changes in the whole larvae, where differential expression in different tissues can confound the interpretation of the results, or in hemocytes, where transcriptional changes reflect mainly hemocyte differentiation [37–40]. Based on these studies, here we investigated the regulation of *Drosophila* humoral anti-parasitoid immunity mediated by the larval fat body. We focused on the *lectin-24A* gene, a key humoral immune effector in this response [12], and studied its induction with parasitization to understand the regulation of anti-parasitoid humoral immune mechanisms more broadly.

## Results

### The upstream cis-regulatory region of the *lectin-24A* gene underlies a specific response to parasitoid wasp infection

We have previously found that the expression of *lectin-24A* increases resistance to parasitoids, and the Drosophila Genetic Reference Panel (DGRP) line 437 carries a resistant allele of the gene whereas the DGRP-892 line carries a susceptible allele [12]. These two alleles differed in their expression levels, where the resistant allele is expressed at a higher level than the susceptible one in homeostasis and when the *lectin-24A* is upregulated after infection [12]. To investigate if the proximal upstream region of the *lectin-24A* gene responds specifically to parasitoid wasp infection, we infected *D. melanogaster* larvae expressing a *lectin-24A* reporter driven by 444 bp of sequence upstream from the transcription start site (TSS) of the resistant DGRP-437 allele (*Venus^LP437^*) [12]. This upstream region was chosen because it is the entire intergenic sequence between *lectin-24A* and its adjacent gene *Shaw*. We infected larvae with virus, bacteria, or parasitoid wasps, and found that only parasitoid wasp infection significantly induces the expression of the reporter (Figure 1A).

**Figure 1.**
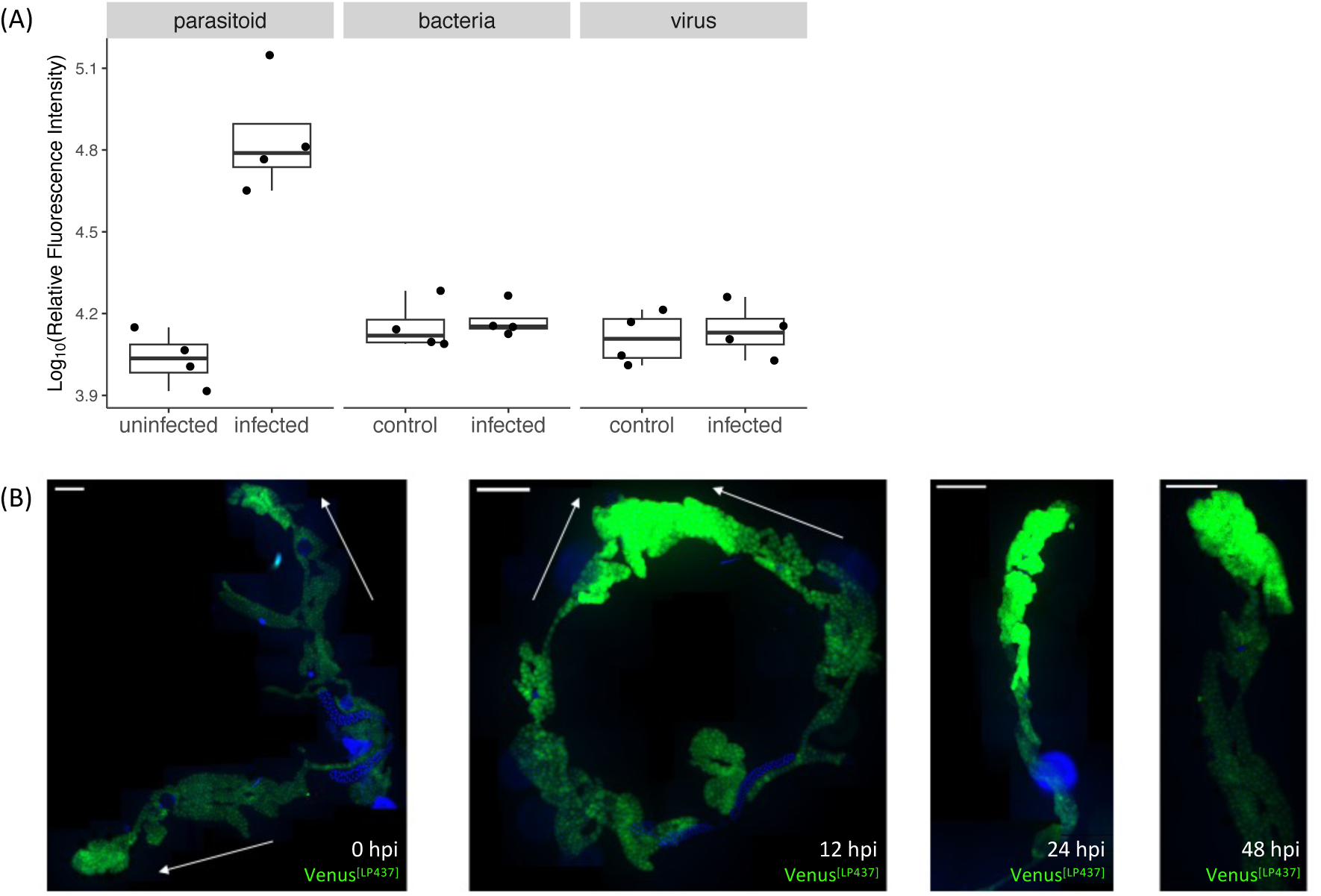
Expression of the *lectin-24A* reporter *Venus^LP437^*. (A) Expression of *Venus^LP437^* 24 hours post-infection (hpi) with different pathogens, left to right: *Leptopilina boulardi* parasitoid wasps, *Escherichia coli* bacteria, and *Drosophila* C virus. The relative fluorescence intensity represents the fluorescence intensity normalized with total protein quantity of each sample. Fluorescence was measured in homogenized larvae with a plate reader in a 96-well plate. Each point represents a pool of 15 larvae. (B) Expression pattern of the reporter *Venus^LP437^* in the *Drosophila* larval fat body. Microscopy imaging of larval fat bodies dissected from uninfected and infected larvae, at 0, 12, 24, and 48 hpi. Fat bodies are shown with posterior pointing up unless indicated by the direction of white arrows. Green represents Venus expression, and blue represents Hoescht nuclear staining. Scale bars represent 500μm.

Consistent with previous reports of *lectin-24A* expression [41], when we observed infected larvae under a fluorescent dissecting microscope the *lectin-24A* reporter *Venus^LP437^* construct was expressed in the larval fat body. We also confirmed the expression by removing the larval fat body through dissection (Figure 1B). Microscopy imaging of the dissected larval fat body shows localization of expression after infection with parasitoid wasps, where the reporter is consistently expressed at higher levels in the posterior ends (Figure 1B).

### RNA sequencing reveals humoral immune response is localized to the posterior larval fat body

As we observed localized expression of *lectin-24A* in the posterior ends of the larval fat body, we investigated whether this localized response is seen in other immune-responsive genes. We conducted fat body RNA sequencing comparing gene expression changes in the anterior and posterior of the fat body in response to parasitoid infection. The fat bodies were separated into anterior and posterior sections according to the red line indicated in Figure 2A. After filtering out lowly expressed genes, we detected a total of 10499 genes expressed in the fat body. Under infected condition, there is a total of 213 significantly differentially expressed genes with absolute log_2_FC of greater than 1, between the anterior and posterior sections. The expression of homeobox (Hox)-containing transcription factor genes, *abd-A* and *Abd-B*, are known for determining the anteroposterior body axis in *Drosophila*, including the fat body, and we see both genes differentially expressed between the anterior and posterior, indicating we have successfully separated the sections by dissection (Figure 2B) [42,43]. The results confirm our observation of localized *Venus^LP437^* expression in the posterior ends of the fat body upon wasp infection, as *lectin-24A* is among the top upregulated genes in the posterior with infection (Figure 2B).

**Figure 2.**
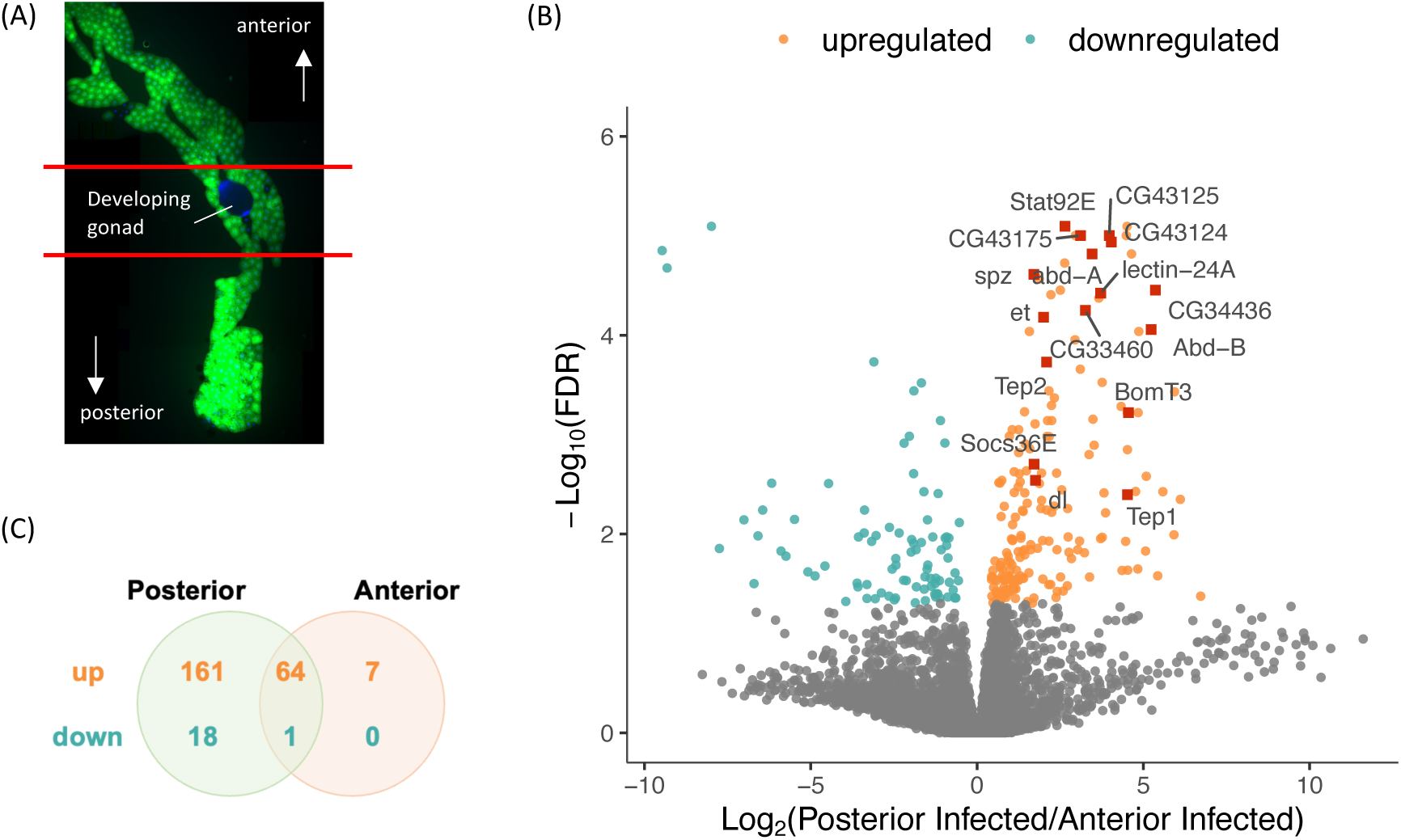
RNA sequencing of larval fat body sections. (A) Dissection of male larval fat bodies along the representative red lines into anterior and posterior sections, where the developing gonads were removed, for RNA sequencing. (B) Differentially expressed genes, with absolute log_2_FC above 1 and False Discovery Rate (FDR) below 0.05, between anterior and posterior larval fat body sections under infected condition. (C) Venn diagram showing overlap of significantly differentially expressed genes with absolute log_2_FC above 1 after infection in anterior and posterior fat body sections.

Moreover, it seems that the posterior of the fat body showed generally higher immune responsiveness than the anterior to parasitoid infection. Following infection, 72 genes were significantly differentially expressed with an absolute log_2_FC of greater than 1 in the anterior, whereas in the posterior the number is 244 genes (Supplementary Figure 2). Most genes differentially expressed in the anterior after infection were also differentially expressed in the posterior, while there were 179 genes that were significantly differentially expressed in the posterior but not in the anterior after parasitization (Figure 2C). We also see that differentially expressed genes in the posterior are enriched for “extracellular” in Cellular Components Gene Ontology (GO) terms, suggesting that the posterior fat body is enriched for secreted proteins (Supplementary Figure 3). In particular, putative immune effectors, including the thioester-containing protein genes *Tep1* and *Tep2*, as well as a number of serine proteases (*CG34436*, *CG33460*, *CG43124*, *CG43125*, and *CG43175*), appear to be specifically upregulated in the posterior of the fat body with parasitization (Figure 2B).

Significantly downregulated genes in the larval fat body after parasitism suggest parasitoid infection is causing shifts in metabolism and other aspects of immunity. *sug* and *daw*, genes involved in sugar metabolism [44,45], are downregulated in the fat body after parasitoid infection. We also found certain immune factors to be downregulated with parasitism, including *Lectin-galC1* and the lysozyme *CG8492*, both involved in antimicrobial response [46,47]. *RhoBTB*, a Rho-related BTB domain-containing protein involved in anti-parasitoid response is downregulated in the posterior fat body after infection [48].

### Key immune signaling pathway components are differentially expressed in the fat body after parasitism

There are also interesting patterns of localized immune signaling gene expressions: key components of the NF-κB and JAK/STAT pathways show up as top differentially expressed genes between posterior and anterior sections in both control and infected conditions (Figure 2B and Supplementary Figure 2). GO enrichment of significantly upregulated genes in the posterior of the fat body also showed enrichment of terms related to “receptor signaling pathway via JAK-STAT”, “immune system process”, and “defense response” (Supplementary Figure 3).

Selecting specific components from major immune signaling pathways including transcription factors, we analyzed their expression patterns in anterior/posterior and uninfected/infected conditions (Figure 3). It is striking to see that different transcription factors show differential patterns of expression. The NF-κB transcription factor *dl* (*dorsal*) is induced specifically in the posterior of the fat body in response to infection, with *Rel* (*Relish*) showing a non-significant trend in the same direction. *Dif* shows similar levels of expression across the four conditions (Figure 3). The gene encoding the Toll receptor *Tl* is also upregulated with infection specifically in the posterior (Figure 3). While the Toll ligand, *spz*, is more highly expressed in the posterior regardless of infection status (Figure 3).

**Figure 3.**
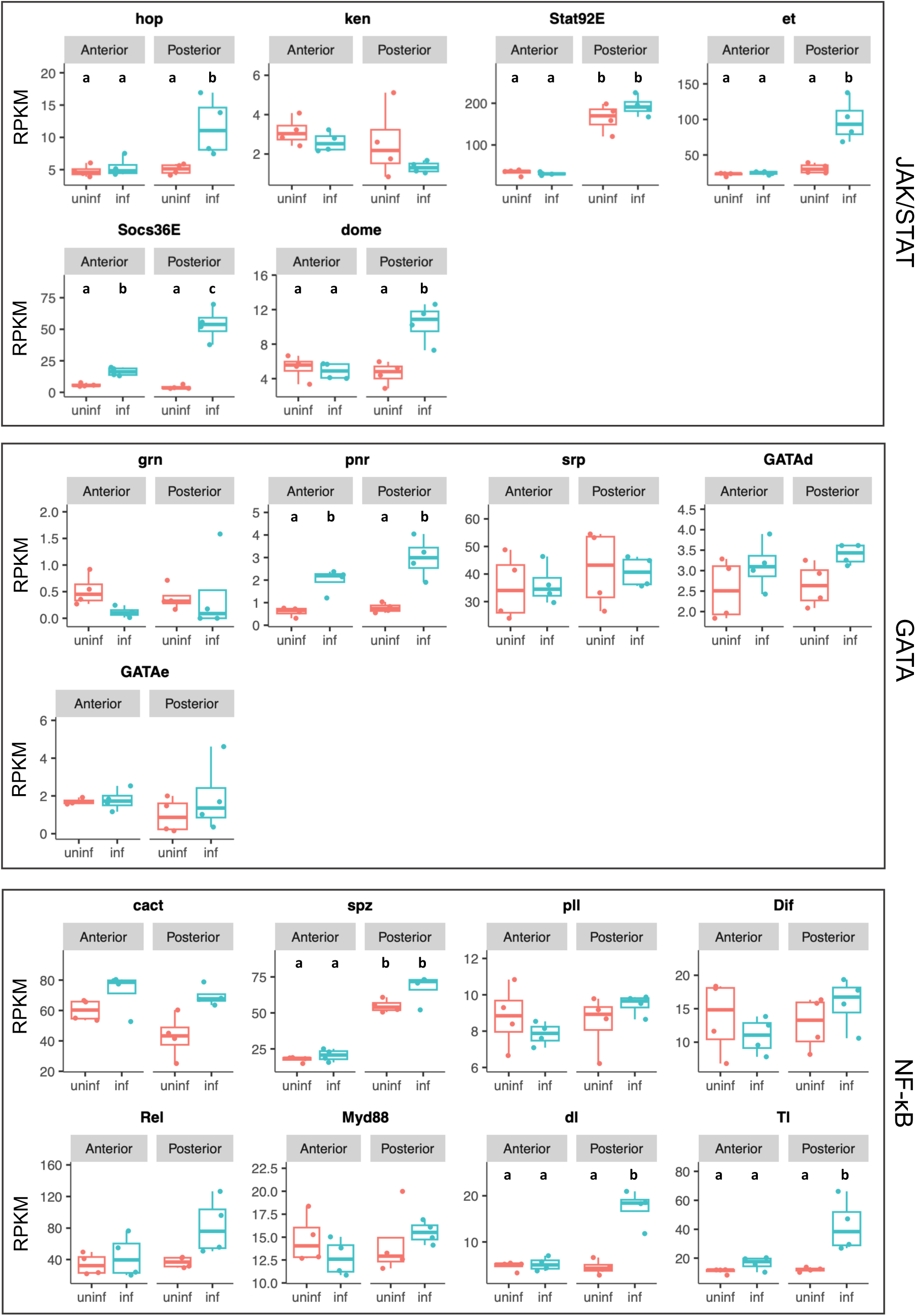
Gene expression patterns of components from major immune pathways in anterior and posterior sections with and without infection. Expression shown as Reads Per Kilobase of transcript per Million mapped reads (RPKM). Labels denote significant (genome-wide false discovery rate < 0.05) differential expression with absolute log_2_FC of at least 1 between treatment groups. Only significant differences are shown.

The five known GATA transcription factors are encoded by *pnr* (*pannier*), *srp* (*serpent*), *grn (grain), GATAd*, and *GATAe*. While the other four GATA transcription factors showed similar expression between the fat body sections and infection statuses, *pnr* is upregulated with infection in both the anterior and posterior (Figure 3).

The JAK/STAT pathway transcription factor *Stat92E* shows an entirely spatial differential expression not affected by infection state, where it is constitutively upregulated in the posterior (Figure 3). However, some other JAK/STAT pathway components have different patterns of expression: *hop* and *dome* are induced specifically in the posterior with infection, while the negative regulator *et* shows the same pattern of posterior induction (Figure 3). Another negative regulator, *Socs36E*, shows induction with infection in both anterior and posterior, but is more strongly induced in the posterior (Figure 3). Overall, we see patterns of localized differential expression of immune pathway component genes in the *Drosophila* larval fat body between the anterior and posterior sections in response to wasp parasitization. The expression patterns of these genes in relation to the localized expression of *lectin-24A* suggest potential roles of these pathways in the regulation of *lectin-24A* and the wider humoral immunity against parasitoid infections.

### Truncation of the upstream promoter sequence of *lectin-24A* defines the minimal region to maintain fat body specific induction of the gene

We have limited understanding of the regulation of the anti-parasitoid immune response in *D. melanogaster*, particularly the early signaling that activates humoral immunity in the larval fat body. We investigated the upstream region of the *lectin-24A* gene to further define the specific sequences that underlie its induction with parasitization, and to understand the transcription factors involved in the regulation of its expression. We created truncated versions of the *Venus^LP437^*reporter construct where we deleted the sequence from the 5’ end (Figure 4A). While we saw little effect from the first two deletions to 420 bp and 314 bp upstream of the gene, we found that, when we truncated the sequence to be 146 bp upstream of the TSS, there were significant changes to the expression of the reporter where it becomes expressed in other tissues such as the gut with and without parasitization (Supplementary Figure 4). Therefore, the region 314 bp upstream of the gene is required for an inducible response of the *lectin-24A* gene that is restricted to the fat body tissue.

**Figure 4.**
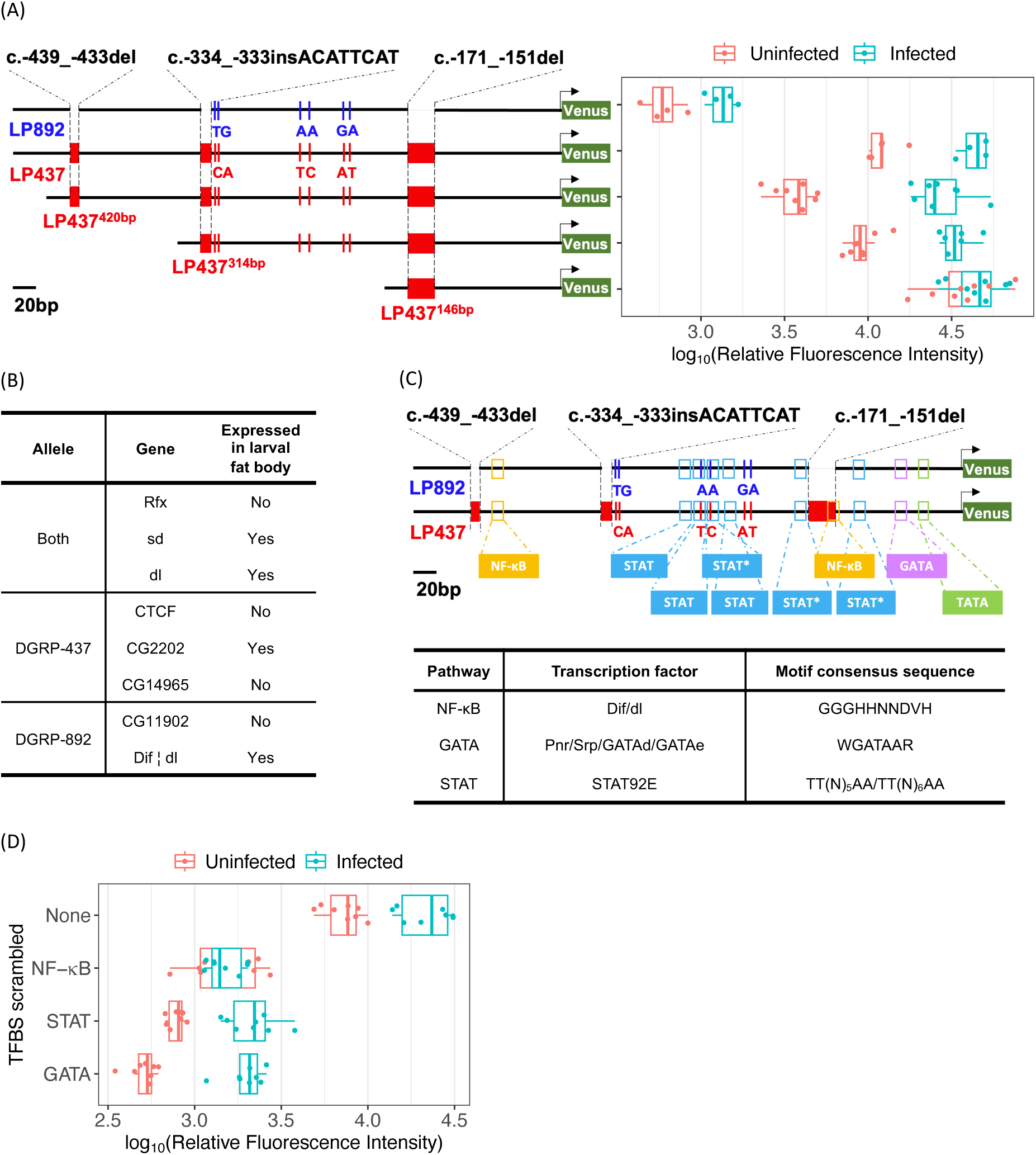
Transcription factor binding motifs (TFBMs) upstream of *lectin-24A*. (A) Truncated *lectin-24A* reporter Venus expression compared to the full-length reporter constructs with the susceptible DGRP-892 (LP892; blue) and resistant DGRP-437 (LP437; red) alleles. (B) Yeast one-hybrid screen results showing proteins that bind to the sequence 444 bp upstream of the resistant DGRP-437 *lectin-24A* allele and/or the susceptible DGRP-892 *lectin-24A* allele. Dif ¦ dl represents a heterodimer. (C) Predicted TFBMs of transcription factors from the JAK/STAT, NF-κB, and GATA pathways upstream of *lectin-24A*. The STAT consensus sequence shown corresponds to the conserved STAT binding site “TT(*n*)AA”, with 5-6 variable *n* sites in the middle. (D) Expression of the reporter constructs with the 314 bp upstream sequence of *lectin-24A* where predicted NF-κB, GATA, and three of the STAT TFBMs are scrambled. The scrambled STAT sites are indicated with asterisks in (C). Each point represents a pool of 10 larvae. The relative fluorescence intensity represents the fluorescence intensity normalized with total protein quantity of each sample.

### NF-κB, GATA, and STAT binding motifs are all involved in the regulation of *lectin-24A*

Our RNAseq results suggest that the NF-κB, GATA, and JAK/STAT pathways may be involved in the regulation of the *lectin-24A* gene and humoral anti-parasitoid immunity in general. We also screened for transcription factors binding to the upstream region of both the resistant and susceptible alleles of *lectin-24A* in a Yeast one-hybrid (Y1H) screen. In particular, we found that dorsal from the Toll pathway binds to both the resistant allele from DGRP-437 and the susceptible allele from DGRP-892 (Figure 4B). We then predicted TFBMs in the 444 bp upstream region according to the consensus binding motifs for transcription factors in each of the immune pathways. One consensus motif was used for the transcription factors Dif/dl, two motifs were used for STAT92E, and four motifs were used for the different GATA transcription factors [49–52] (Figure 4C & Supplementary Table 1). We predicted TFBMs for each of these transcription factors in this region (Figure 4C). Furthermore, the GATA motif is predicted to bind all four of the GATA transcription factors (Supplementary Table 1).

To further study the role of TFBMs in the regulation of *lectin-24A*, we created promoter reporter constructs where motifs have been scrambled to abolish transcription factor binding. Since the truncated version with 314 bp upstream of the TSS showed similar expression as the full-length LP437 construct, we scrambled the TFBMs with this version of the reporter (Figure 4A, LP437^314bp^). The universally conserved STAT binding site is defined as “TT(*n*)AA”, with variable 4-6 *n* sites in the middle [53,54]. The *Drosophila* STAT binds “TTC(N)_3_GAA” sites with high affinity [50,51]. However, gene regulation can rely on STAT binding motifs that have lower affinity [50]. This includes *Dome,* which is upregulated by STAT binding “TTC(N)_4_GAA” sites [50], and which we found to be upregulated in the posterior fat body after infection (Figure 3). Here we scrambled three sites with the motif “TTC(N)_5_AA” in the *lectin-24A* upstream region, which are half-way between the universal STAT motif and the 4*n Drosophila* STAT motif (Figure 4C, sites with asterisks & Supplementary Table 2). We also scrambled motifs predicted to bind Dif/dl and GATA.

We found that the predicted TFBMs associated with Dif/dl, STAT92E, and GATA are all involved in the regulation of the gene. Scrambling of the TFBM for Dif/dl resulted in the abolishment of the upregulation of the reporter and reduction in the basal expression, whereas scrambling of the TFBMs for the GATA and STAT transcription factors resulted only in the reduction of the basal expression while the induction with infection is still present (Figure 4D). These results suggest that all three of these pathways may bind to the upstream region of *lectin-24A* and be involved in the regulation of expression.

We previously found that the *cis*-regulatory sequences of the resistant and susceptible alleles of *lectin-24A* differed by three insertion-deletion polymorphisms (indels) of 7 bp, 8 bp, and 21 bp in length, and six single nucleotide polymorphisms (SNPs; Figure 4A, top two rows) [12]. The 21 bp indel was found to be primarily responsible for the differential regulation of *lectin-24A* expression between the two alleles [12]. Based on the positions of the TFBMs in the region, we found that the NF-κB binding site is disrupted by the 21 bp deletion in the susceptible DGRP-892 *lectin-24A* allele (Figure 4C). As scrambling this NF-κB binding site abolishes the basal expression and induction of the reporter, we conclude that the loss of this NF-κB binding site explains the differential expression we see from natural polymorphisms in the *lectin-24A* upstream region. This in turn controls susceptibility to infection in natural populations [12].

### The JAK/STAT pathway plays a crucial role in the regulation of *lectin-24A*

The JAK/STAT signaling pathway is an evolutionarily conserved pathway with a variety of functions in hematopoiesis, immunity, and development [55]. The canonical JAK/STAT signaling starts with ligand binding, resulting in receptor dimerization bringing two non-receptor tyrosine kinases (JAKs) together to trans-phosphorylate one another [56,57]. Activated JAKs then phosphorylate the receptor which in turn phosphorylates the STAT transcription factors. This leads to the normally cytosolic STAT molecule translocating to the nucleus to activate gene expression. The *Drosophila* JAK/STAT pathway only has one JAK (*hop*) [58], one active type I cytokine receptor (*dome*) [59], and one STAT (*Stat92E*) [51,60]. To better characterize the role of the JAK/STAT pathway in the regulation of *lectin-24A*, we knocked down the expression of the transcription factor Stat92E in larvae expressing the *lectin-24A* reporter using the ubiquitously expressed *da-GAL4* to drive the expression of a hairpin RNAi construct. At 24 hpi, *Stat92E* knockdown abolished the induction of the reporter, suggesting that the JAK/STAT pathway is involved in the regulation of the gene (Figure 5A).

**Figure 5.**
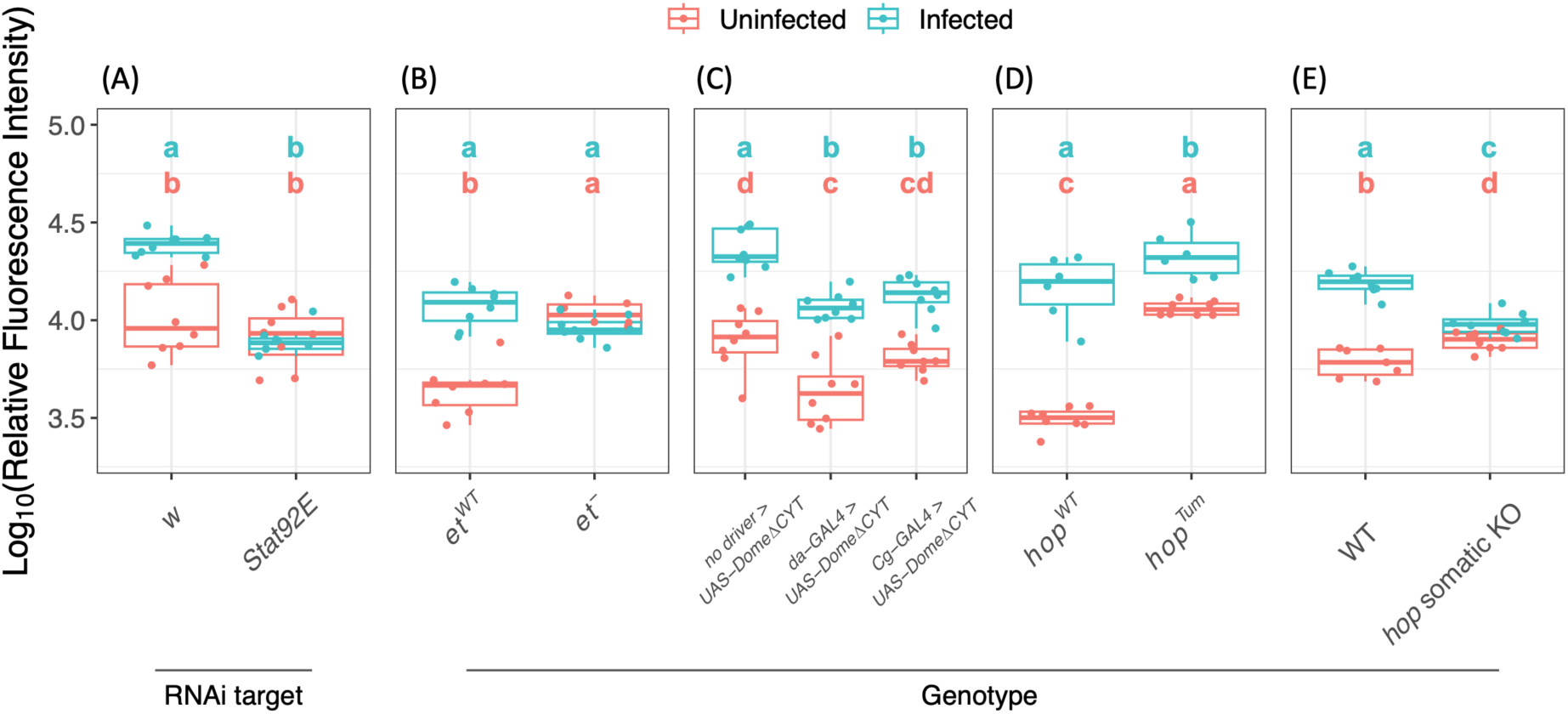
The role of the JAK/STAT pathway in the regulation of *lectin-24A*. In all panels the *Venus^LP437^* reporter expression was measured. (A) Larvae with knockdown of *Stat92E* using RNAi, compared to the knockdown of the *white* gene as a control. (B) Larvae carrying a mutant or wildtype (WT) form of the JAK/STAT negative regulator *et*. (C) Larvae expressing a dominant negative form of the JAK/STAT receptor Dome (DomeΔCYT) in the fat body and hemocytes under the control of the *Cg-GAL4* driver, with the ubiquitous *da-GAL4* driver, and with no driver. As the parental stock expressing *UAS-DomeΔCYT* also expresses the JAK/STAT negative regulator *et* under UAS control on a balanced chromosome II, a portion of the larvae also expresses *UAS-et*. Larvae expressing GAL4 drivers also expressed *tub-GAL80^ts^*. (D) Larvae expressing the constitutively active *hop* (*hop^Tum^*) compared to larvae expressing the wild type (WT) *hop*. (E) Larvae expressing *Act-Cas9* and guide RNA targeting *hop* compared to control larvae. Each point represents a pool of 10 larvae. The relative fluorescence intensity represents the fluorescence intensity normalized with total protein quantity of each sample. All infected larvae were collected at 24 hpi and uninfected age-matched larvae were collected for the controls. Tukey’s test for each panel, *P* < 0.05 between groups.

In our RNAseq analysis, we also found that the JAK/STAT pathway negative regulator *et* showed specific induction in the posterior of the fat body. et is structurally similar to the JAK/STAT receptor dome, and it can function as a negative regulator by forming inactive heterodimers with dome [21,61]. It was previously found that et is required for the downregulation of JAK/STAT activity in hematopoietic progenitors for lamellocyte differentiation after wasp infection [21]. To further understand the role of JAK/STAT signaling in *lectin-24A* regulation, we analyzed the expression of the reporter in a genetic background where *et* is mutated [21]. We found that with a non-functional mutant *et*, the reporter shows constitutively upregulated expression in the larval fat body (Figure 5B). In addition, to confirm this result, we used a dominant negative allele of Domeless (DomeΔCYT, cytoplasmic domain deleted), driven in the fat body by *Cg-GAL4* or ubiquitously by *da-GAL4*, to decrease JAK/STAT activity. We observed a general lowering of reporter expression both for basal and infected conditions (Figure 5C).

The *Drosophila* JAK, *hop*, showed a similar expression pattern as *et* in our RNAseq analysis, being upregulated after infection in the posterior fat body. We therefore studied the expression of the reporter in a genetic background with a constitutively active form of *hop* (*hop^Tum^*). As predicted, the reporter showed elevated basal expression in the *hop^Tum^* background (Figure 5D). When we then examined *lectin-24A* reporter expression in a background where we abolished *hop* in somatic cells using CRISPR/Cas9 and a guide RNA targeting the gene (somatic knock out), we found a reduction in the upregulation of the reporter after wasp infection (Figure 5E). Taken together, our results indicate clearly that the JAK/STAT pathway has a critical role in the induction of *lectin-24A* following parasitization.

### GATA transcription factors modulate *lectin-24A* expression in the fat body

GATA transcription factors comprise a family of DNA binding proteins named after the consensus sequence (WGATAAR) they recognize. GATA factors are highly conserved across invertebrates and vertebrates, where they play major roles in development. *Drosophila* and mammalian GATA factors share functional similarities in the regulation of developmental processes, such as hematopoietic development [26], blood cell differentiation [28,62], and liver/fat body development [28,63,64]. *Drosophila* has a total of five GATA factors, of which only three display the canonical two zinc finger domains found in all mammalian GATA factors: Pannier (Pnr), Serpent (Srp) and Grain [65–67].

The DNA-binding activity of GATA factors is mainly mediated by the C-terminal zinc finger domain, whereas the N-terminal zinc finger domain is required for interactions between the GATA factor and its cofactor Friend of GATA (FOG) proteins [68]. It has previously been shown that the FOG protein u-shaped (Ush) interacts with the N-terminal zinc finger of both Pnr and Srp [65,68,69]. In particular, Pnr and Ush form heterodimers, where Ush antagonizes the activator function of the Pnr transcription factor [65,69,70].

As we observed significant differential expression of *pnr* in our RNAseq analysis following parasitization (Figure 3), we studied the effect of Pnr on *lectin-24A* expression. We utilized the *pnr^D4^* mutant, which belongs to a class of *pnr* mutants with point mutations in the N-terminal zinc finger domain, resulting in very weak interactions with Ush [65,69]. Using the fat body *Cg-GAL4* driver, we expressed the dominant *pnr^D4^* in the fat body with the *UAS-pnr^D4^* construct (Figure 6A). As *pnr^D4^* interacts poorly with the negative Pnr regulator Ush, it functions as a dominant positive mutant. Strikingly, *Venus^LP437^* reporter showed constitutive high expression, even in the absence of parasitization (Figure 6A). We then analyzed the expression of *lectin-24A* Venus reporter in a genetic background where we knocked out *pnr* expression using somatic mutagenesis with CRISPR/Cas9. However, knock out of *pnr* expression in the whole larvae did not result in significant changes to the *Venus^LP437^* reporter expression, which might reflect GATA factors may have redundant roles in regulating *lectin-24A* (Figure 6B). Thus, our results indicate that Pnr is involved in the induction of *lectin-24A* following parasitoid infection, but it may not be the only or the main GATA factor playing this role.

**Figure 6.**
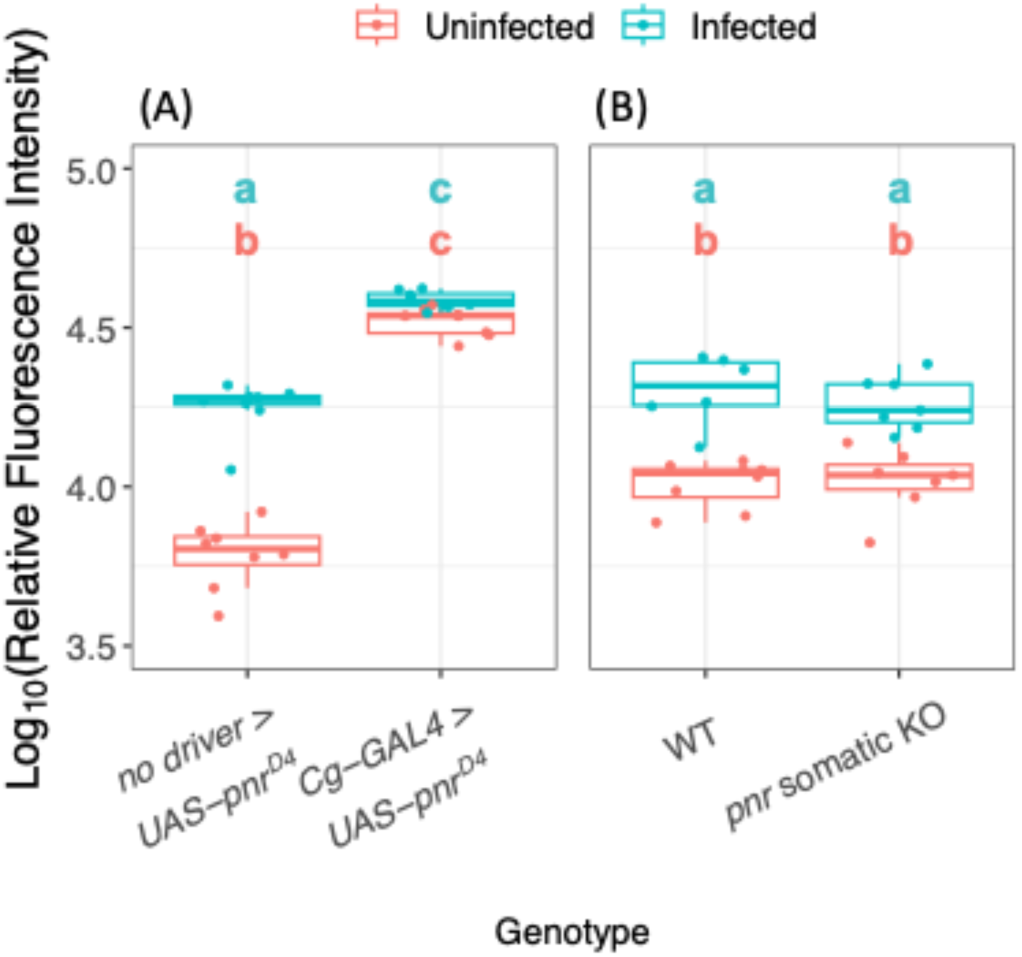
The role of the GATA transcription factor, Pannier, in the regulation of *lectin-24A*. The *Venus^LP437^* reporter expression was measured. (A) Larvae expressing the dominant positive form of Pnr (Pnr^D4^) under UAS control driven by the fat body and hemocyte-specific *Cg-GAL4* driver or no driver. (B) Larvae expressing *Act-Cas9* and guide RNA targeting *pnr* compared to control larvae. Each point represents a pool of 10 larvae. The relative fluorescence intensity represents the fluorescence intensity normalized with the total protein quantity of each sample. Tukey’s test for each panel, *P* < 0.05 between groups.

### NF-κB transcription factors dl regulates humoral antiparasitoid immunity

NF-κB transcription factors of the Toll and Imd pathways have major roles in the regulation of antimicrobial humoral immunity for *Drosophila*. Our scrambling of TFBMs in the *lectin-24A* upstream region suggests that the Toll pathway might also participate in regulating the anti-parasitoid response in the larval fat body (Figure 4D). Of the three *Drosophila* NF-κB transcription factors, Dif, dl, and Rel, we found that only dl becomes significantly upregulated with parasitism specifically in the posterior of the larval fat body (Figure 3). To further understand the role of NF-κB transcription factors in the regulation of *lectin-24A*, we created somatic mutants of the NF-κB transcription factors Dif and dl. The effects of somatic mutagenesis of these genes suggest that knocking out *dl* reduces the expression of *Venus^LP437^*in the uninfected state (Figure7A), while removing *Dif* expression seems to give the opposite effect (Figure 7B). We then overexpressed dl and Rel in the larval fat body using the driver *Cg-GAL4*, where we show that dl overexpression leads to a constitutively high expression of the *Venus^LP437^* reporter with and without infection (Figure 7C). Overexpression of Rel did not show significant effects on the reporter expression (Figure 7C). Altogether, we conclude that dl is the NF-κB transcription factor regulating *lectin-24A* expression.

**Figure 7.**
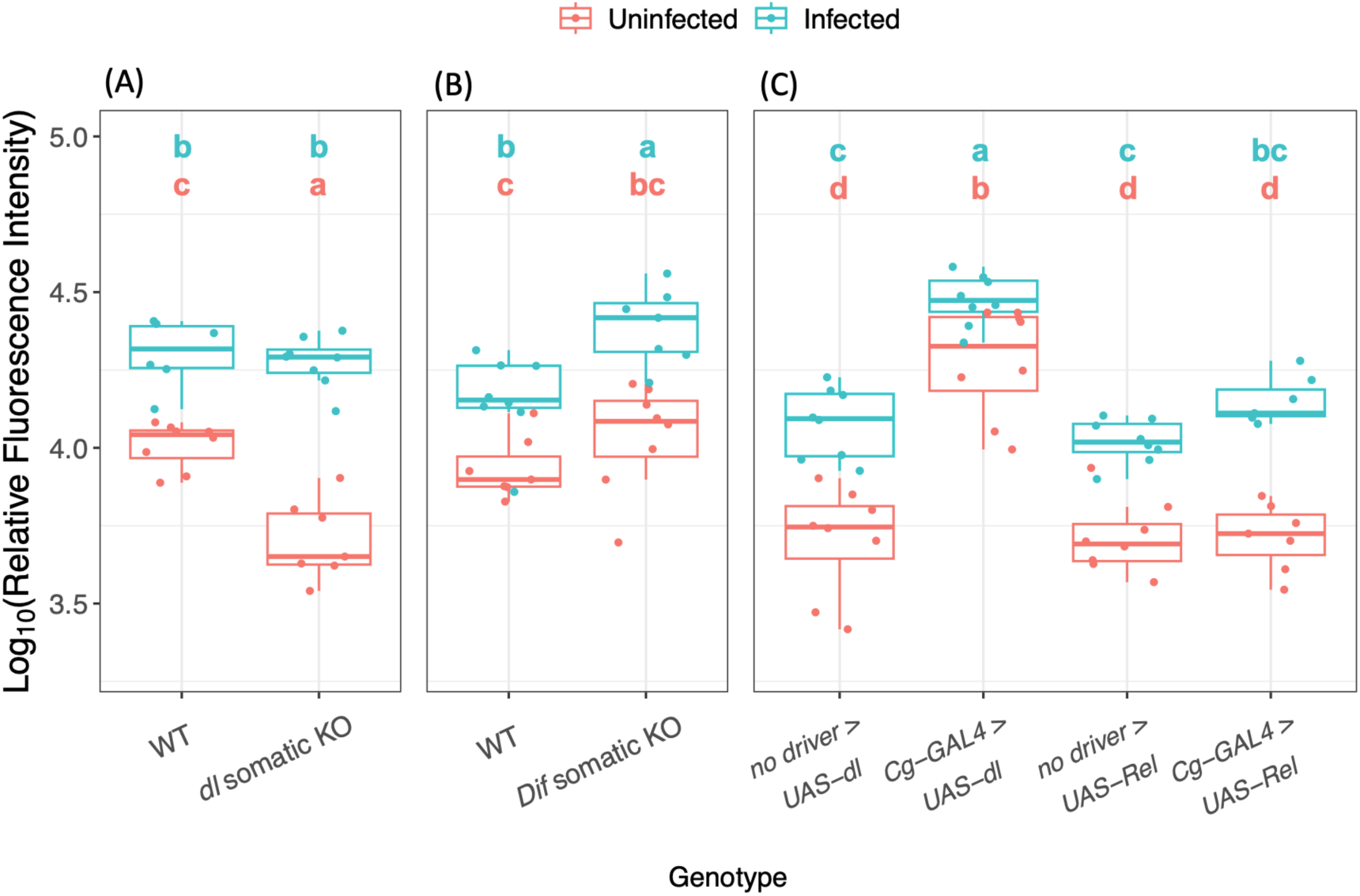
The role of NF-κB transcription factors in the regulation of *lectin-24A*. The *Venus^LP437^*reporter expression was measured. (A) Larvae expressing *Act-Cas9* and guide RNA targeting *dl* compared to control larvae. The same WT control samples were used here as in Figure 6B. (B) Larvae expressing *Act-Cas9* and guide RNA targeting *Dif* compared to control larvae. (C) Larvae expressing *dl* or *Rel* under UAS control driven by the fat body and hemocyte-specific *Cg-GAL4* driver or no driver. F_1_ heterozygote larvae were collected. As the parental line expressing *UAS-dl* carried a balancer chromosome that could not be identified at the larval stage in F_1_ progeny, larvae collected from the *UAS-dl* crosses were a mix of *Cg-GAL4/UAS-dl* and *Cg-GAL4/+*. Each point represents a pool of 10 larvae. The relative fluorescence intensity represents the fluorescence intensity normalized with the total protein quantity of each sample. Tukey’s test for each panel, *P* < 0.05 between groups.

### Transcription factor binding motifs found in the upstream region of *lectin-24A* are generally enriched in the upstream sequences of parasitism-induced genes

As we have found the TFBMs of GATA, NF-κB, and STAT are important in the regulation of the *lectin-24A* gene, we were interested to find out if this generalizes to the other genes upregulated with infection. We searched for TFBMs in the 300 bp upstream region of all the genes detected in the larval fat body RNAseq (Figure 2). Then we assessed if a TFBM is enriched in the upstream sequence of genes upregulated with infection in the posterior of the fat body, relative to the full set of genes detected. We found that GATA binding motifs are generally enriched around two-fold in the sequences upstream of upregulated genes. For instance, the Pannier binding motif (Supplementary Table 1) is found an average of 0.60 times per kilobase (kb) of sequence 5’ of TSSs for upregulated genes, whereas it is only found an average of 0.33 times per kb of upstream sequences in the full set of genes (Welch’s *t*-test, t = 3.49, d.f. = 277.9, *p*-value = 5.65 × 10^-4^). A similar case is observed in the genes specifically upregulated in the posterior of the fat body after infection (Figure 2B), showing even higher enrichment with a mean of 0.66 times per kb in these genes (Welch’s *t*-test, t = 3.51, d.f. = 186.1, *p*-value = 5.72 × 10^-4^).

When we searched for STAT motifs in the upstream sequences, we found significant enrichment of the “TTC(N)_4_AA” motif in the posterior-specific upregulated genes with a mean of 8.46 instances per kb compared to 7.77 in the background (Welch’s *t*-test, t = 2.10, d.f. = 188.84, *p*-value = 0.037). Although the same motif also appears to be enriched in the parasitism-induced gene set, with 8.27 occurrences per kb, it falls just below the significance threshold (Welch’s *t*-test, t = 1.91, d.f. = 284.98, *p*-value = 0.058). On the other hand, the “TTC(N)_5_AA” motif only shows significant enrichment in the parasitism-induced gene set, with 6.29 occurrences per kb compared to 5.83 in the background set (Welch’s *t*-test, t = 1.98, d.f. = 284.47, *p*-value = 0.048).

Moreover, previous studies have described a specific organization of TFBMs in the upstream region of fat body-specific immune genes, where NF-κB and GATA binding sites have been found to be in close association, with less than 50 bp in-between them and within 200-300 bp upstream of the TSSs [30,31]. We specifically searched for NF-κB sites within 50 bp of a GATA site and checked for enrichment. Although, we see that NF-κB motifs linked to GATA motifs are also more highly enriched in the upstream region of parasitism-induced genes and posterior-specific anti-parasitoid genes, with a mean of 0.52 and 0.53 instances per kb, respectively, compared to 0.39 in the background set, the differences were not significant. Overall, the TFBMs of the GATA, NF-κB, and STAT transcription factors all appear to be enriched in the 5’ sequences of parasitism-induced genes in the *Drosophila* larval fat body, suggesting that these transcription factors and the signaling pathways they belong to are generally involved in the regulation of genes in the humoral anti-parasitoid response.

## Discussion

The *Drosophila* anti-parasitoid immune response involves the concerted action of hemocytes and the fat body to encapsulate and melanize the wasp [4,63,64]. This complex and multifaceted response is distinct from responses to microbial or viral infection, and is specifically induced by a wasp PAMP [32]. Here we have demonstrated that the regulation of *Drosophila* anti-parasitoid humoral immunity, mediated by the larval fat body, involves complex inputs from multiple immune pathways (Figure 8).

**Figure 8.**
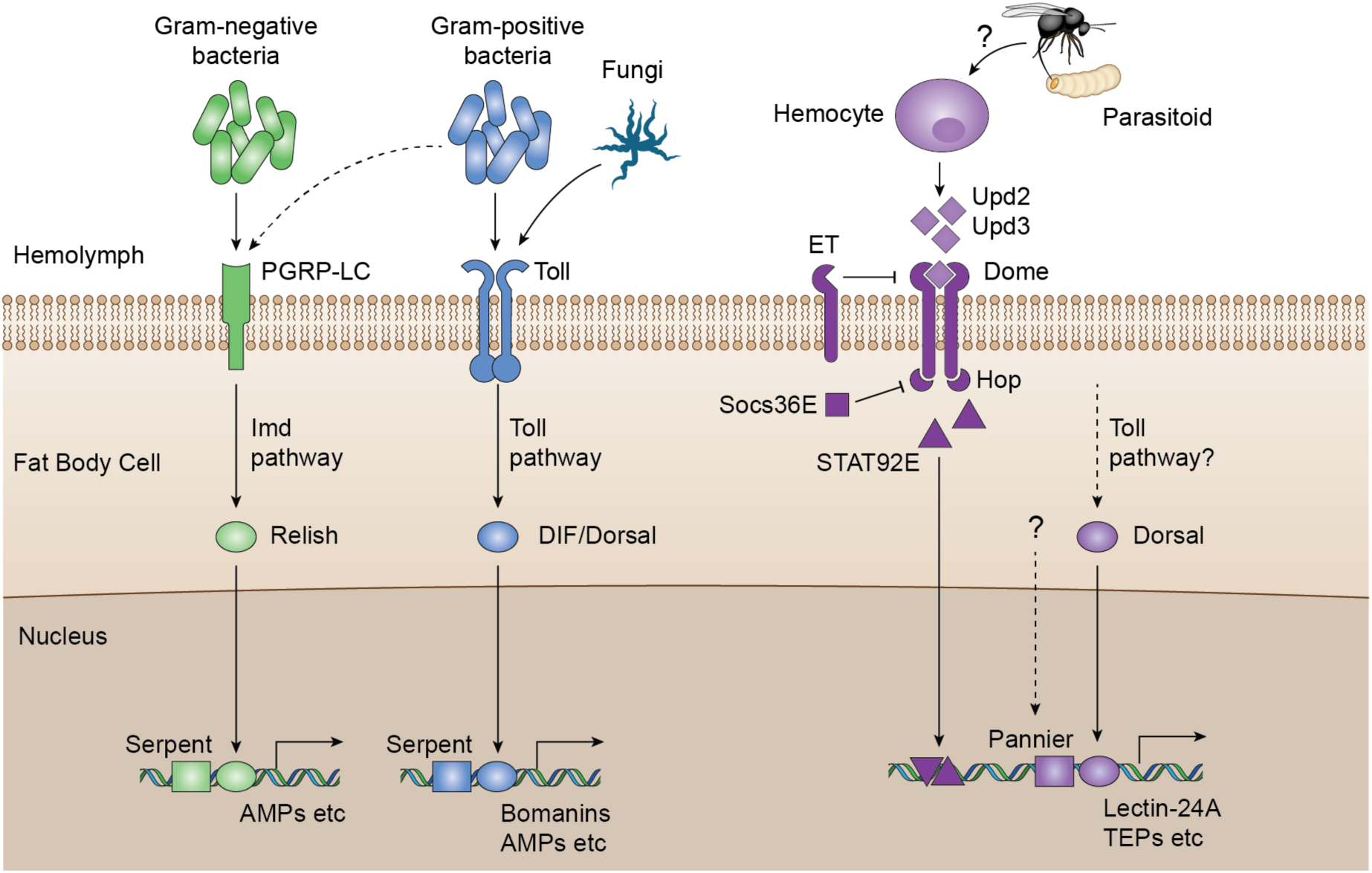
Schematic overview of *Drosophila* humoral defenses to microbial pathogens and parasitoid. Antimicrobial responses in the fat body are regulated by the NF-κB pathways, Toll and Imd. The Imd pathway is mostly activated by Gram-negative bacteria through the PGRP-LC receptor, whereas the Toll pathway is largely activated by fungi and Gram-positive bacteria through the Toll receptor. Activation of the Imd pathway leads to the NF-κB transcription factor Relish translocating into the nucleus after its cleavage and activation, and the transcription factors Dif and Dorsal are activated through Toll pathway signaling. These NF-κB transcription factors along with the GATA transcription factor Serpent induce the expression of genes encoding antimicrobial effectors in the fat body via closely linked GATA and κB binding sites in the upstream promoter region. The *Drosophila* humoral anti-parasitoid response is regulated by a combination of inputs from JAK/STAT, GATA and NF-κB pathways. After recognition of parasitism, hemocytes produce the cytokines, Upd2 and Upd3, which bind to the receptor Dome to activate JAK/STAT pathway signaling in the fat body. Dome dimerization after ligand binding leads to trans-phosphorylation of Hop, which then activates the transcription factor STAT92E. STAT92E binds the upstream region of anti-parasitoid genes including *lectin-24A*. ET and SOCS36E are negative regulators of the JAK/STAT pathway that suppress this signaling. STAT, GATA, and NF-κB TFBMs are found to be enriched in the upstream sequences of parasitism-induced genes. The GATA transcription factor Pannier and the NF-κB transcription factor Dorsal also participate in the regulation of anti-parasitoid genes in the larval fat body by binding to their respective binding sites in the promoter region.

The transcriptional response induced by a parasitoid attack differs from that activated by microbial infections [33]. We found that *lectin-24A* is a gene specifically induced by parasitoid infection, providing a tool to understand broadly how wasp-responsive genes are regulated. Based on our current results, it seems that this specificity is potentially conferred by the coordinated inputs from different immune-signaling pathways: NF-κB, the GATA factor Pannier, and JAK/STAT. This regulation differs from the well-characterized antimicrobial defense, which mainly requires the two NF-κB pathways, Toll and Imd, with additional inputs from the GATA transcription factor Serpent [30,74–78].

It is likely that our results reflect mechanisms of regulation in the wider humoral anti-parasitoid response, not just limited to *lectin-24A*. Our analysis of upstream regulatory motifs found enrichment of NF-κB, GATA, and STAT binding sites in the upstream region of differentially expressed genes in the larval fat body in response to wasp infection. This result is consistent with a previous report of TFBMs significantly associated with anti-parasitoid genes in the whole larvae [33]. In that study, Wertheim et al. defined clusters of differentially expressed genes based on patterns of co-expression over nine time points in the 72 hours after parasitization [33]. *lectin-24A* as well as the genes encoding three thioester-containing proteins (TEPs), which are structurally similar to mammalian complement proteins, were clustered together [33]. Genes in this cluster showed significant over-representation of NF-κB, GATA, and STAT regulatory motifs [33], supporting out conclusion that the anti-parasitoid humoral immune response mediated by the fat body is regulated by the concerted action of these pathways. Further support comes from analyses of TEP expression in flies where signaling pathways have been perturbed. TEPs are expressed mainly by the *Drosophila* fat body and induced in response to various immune challenges [79]. The loss of the four secreted TEPs, TEP1-4, results in reduced capacity to successfully encapsulate wasps [8]. *Tep1* shows constitutive expression in the absence of infection in the constitutively active *hop^Tum^* mutant [79]. Similarly, in a *Toll* gain-of-function mutant background, *Tep1* also displayed constitutive activation [79]. However, in this case, the Toll pathway effects were recognized to be indirect, and secreted TEPs were found to function upstream of Toll and promote Toll activation [8,79].

It is still unknown how parasitoids are recognized by the *Drosophila* host. As hemocytes are responsible for the secretion of Upd cytokines that activate the JAK/STAT pathway in other tissues, they are likely to play a key role in the recognition of parasitoid attacks [22]. Our study shows a significant role of the JAK/STAT pathway in *lectin-24A* regulation. Interestingly, we see that JAK/STAT pathway negative regulators are strongly induced alongside the activation of the pathway itself (Figure 3) [21,61,80,81]. As we only investigated one time point (24 hpi) in our experiments, this result suggests potential negative feedback loops in the shutdown of the pathway that were not captured. In mammals, negative regulators of the pathway are known to be induced after infection and JAK/STAT activation, playing a key role in regulating the cytokine response and preventing damaging inflammation [82,83]. This is also the case in *Drosophila*, where expression of the negative regulator SOCS36E is activated by JAK/STAT signaling, creating feedback inhibition [81,84,85].

In the *lectin-24A* upstream region, we predicted a GATA transcription factor binding site that contains the consensus binding motifs from four of the *Drosophila* GATA transcription factors (Figure 4 & Supplementary Table 1). We specifically focused on Pannier because it is the only GATA transcription factor showing differential expression in the larval fat body following parasitoid attack (Figure 3). Our results indicate that Pannier is involved in regulating *lectin-24A* and likely the broader humoral anti-parasitoid response. However, we have yet to investigate whether upregulation of Pannier expression is a primary signal of infection or a consequence of activating other pathways such as JAK/STAT. Furthermore, whether the other *Drosophila* GATA transcription factors also contribute to regulation remains to be further studied. In particular, Serpent, which is known to be involved in the regulation of immune genes in the larval fat body, is also predicted to bind upstream of *lectin-24A* and might also have a role in the regulation of immunity against parasitoids [30,31,86].

Based on our RNAseq data, components of the Toll pathway showed significant differential expression in the larval fat body following parasitism (Figure 3). Particularly, of the NF-κB transcription factors, *dl* showed upregulation specifically in the posterior fat body with infection (Figure 3). We also found that a Dif/dl binding motif in the *lectin-24A* upstream region had a crucial role in the expression of the *Venus^LP437^* reporter (Figure 4D). Dif and dorsal, the two transcription factors in the *Drosophila* NF-κB Toll pathway, are known to regulate the defense to Gram-positive bacteria and fungi. In the antimicrobial response, Dif and dl have different roles depending on developmental stage. In adults, Dif alone is required for the humoral immune response, whereas in larvae, it shows some redundancy with dl [76,87]. While there is evidence that in Dif and dl double mutants, expression of either transcription factors is sufficient to rescue the expression of AMPs such as *Drosomycin* [87], some other reports show that induction of certain AMPs specifically require Dif and cannot be rescued with only dl [77]. Furthermore, Dif and dl, as well as Rel, form homodimers and heterodimers during DNA binding, meaning that permutations of different dimers expand the regulatory possibilities from these NF-κB proteins [88,89]. Since NF-kB transcription factors activate transcription by translocating into the cell nucleus, our observation that the expression of *Dif* does not increase after infection does not preclude it playing a role in regulating the anti-parasitoid humoral response. Nevertheless, as *dl* but not *Dif* mutants reduced expression of the *Venus^LP437^* reporter, dl is likely playing the key role in regulation of this immune response.

In this study, we investigated individually how each of these pathways contributed to the regulation of *lectin-24A* expression, but it remains unclear how they work together. It was previously found that NF-κB and GATA regulatory synergy plays a key role in many immunity-related fat body-specific genes that are activated in response to microbial infections [30,31]. In particular, a fixed organization of NF-κB and GATA binding sites in the 5’ regulatory region within 1 kb of predicted transcription start sites (TSSs) was defined, where both sites are located in the proximal region within 200 or 300 bp from the TSS [30]. The sites were found to be closely linked, within 50 bp from each other and in the same relative orientation [30]. The NF-κB and GATA binding sites we identified in the *lectin-24A* upstream region fit this criterion exactly (Figure 4C). According to the previous study, each individual NF-κB and GATA site has an essential role in the optimal expression of the genes [30]. Furthermore, their results suggest that the GATA site is responsible for localizing the expression to the fat body, and the NF-κB site is responsible for the induction of the gene in response to infection [30]. Our analysis based on scrambling the binding sites in the *lectin-24A* reporter construct are also in line with this model, with the addition that STAT binding is also involved in regulation in a similar way as GATA (Figure 4D). However, notably, Pannier is strongly upregulated by parasitoid infection and this upregulation does not differ between anterior and posterior of the larval fat body (Figure 3). This result suggests that Pannier is not just playing a role to localize *lectin-24A* expression to the fat body, but is also involved in the induction of its expression. Nonetheless, when we manipulated specific genes in each of these pathways the effects on *lectin-24A* expression become less straightforward, suggesting more complex models of coordinated regulation and potential indirect effects from knockdown or knockout in the whole larva.

Immune systems evolve rapidly as hosts adapt to an ever-changing array of parasites and pathogens. We have previously shown that a polymorphic deletion upstream of Lectin-24A has led to reduced parasitoid resistance in wild populations [12], and here we provide an explanation of this by showing that it abolishes an NF-κB binding motif. Furthermore, a group of four single nucleotide polymorphisms cause a smaller reduction in Lectin-24A expression [12]. Notably, one of these is within a STAT binding motif, with the low expression allele predicted to bind STAT92E with lower affinity (TTC(N)_4_AA in the high expression DGRP-437 line becomes TTA(N)_4_AA in the low expression DGRP-892 line). Therefore, the cis-regulatory elements that respond to immune signaling pathways may be hotspots of host-parasite coevolution.

## Methods

### Lectin promoter reporter assay with bacterial, viral, or wasp infection

Six vials of cornmeal fly food (per 1200 ml water: 13g agar, 105g dextrose, 105g maize, 23g yeast, 35ml Nipagin 10% w/v) were prepared for overnight egg lay with the LP437 line, where three vials were used for wasp infection and three vials used as uninfected controls. Three hundred flies of the LP437 line, with approximately even distribution of males and females, were prepared in a cage for egg lay onto a 90mm agar plate coated with a thin layer of yeast paste. After an overnight egg lay in the cage onto an agar plate, the eggs were transferred onto 50mm plates containing cornmeal fly food and some yeast. The eggs were removed from the agar plate by washing them down with a phosphate-buffered saline (PBS) solution (prepared from tablets Oxoid# BR0024G) and dislodging the eggs with a clean plastic brush. The egg solution was transferred into a 15ml Falcon tube and fly eggs allowed to sink to the bottom, where 500μl were then transferred using a cut pipette tip into an Eppendorf tube. Using a cut 200μl pipette tip, 15μl of fly eggs were transferred onto each 50mm plate. Four plates were prepared. The vials and plates were all incubated at 25°C and 70% humidity, and on the fourth day infections were carried out: three female parasitoid wasps were added to each of the three vials for infection for three hours; bacterial and viral infections were carried out according to the protocol below. At 24 hours post-infection (hpi), larvae were collected from the fly food by adding 15% sugar solution to allow the larvae to float to the top of the vial. Groups of fifteen larvae were collected and transferred to a piece of filter paper to dry. They were then transferred to a PCR tube containing 5-10 1.0mm zirconia beads (BioSpec Products # 11079110zx) with 100μl PBS (Gibco# 10010-015). This procedure was repeated until we have four samples for each infected and uninfected conditions with each pathogen.

### Bacterial and viral cultures for infection

A LB-agar plate with no antibiotic added was prepared under sterile conditions, *E.coli* of the OP50 strain was transferred to the plate using a sterile inoculation loop. The loop was then wiped around the plate to produce single colonies. One colony was picked from the plate following an overnight incubation at 37°C. The colony was then transferred to 5ml of sterile LB with no antibiotic. The culture was shaken overnight at 37°C. We then recorded the initial concentration of the *E. coli* culture at OD600 (600nm) by measuring absorbance with a NanoDrop^TM^. 1ml of the bacterial culture was transferred to a 1.5ml Eppendorf tube and spun down at 10,000xg for 5 minutes to concentrate the culture. It was then diluted according to the initial concentration to be slightly above 0.5 at OD600. This culture was then read again on the nanodrop and diluted further to be exactly 0.5 at OD600. The DCV viral cultures were prepared according to Bruner-Montero et al., and aliquots were thawed immediately before use [90].

### *Drosophila* larval infection with virus and bacteria using tungsten wires

We cut a short section of about 1-2 cm of Tungsten wire using scissors or a wire cutter. The wires were mounted into nickel plated holders and adjusted to an appropriate length. The tips of the Tungsten wires were sharpened by connecting them to a circuit with Sodium Hydroxide (NaOH) solution and a battery pack. While sharpening, we checked the tips of the Tungsten wires continuously to see if it needs to be sharpened further.

We transferred the prepared bacterial or viral culture to the caps cut off from 0.5ml Eppendorf tubes and the caps were filled as much as possible. Control cultures were also prepared for either bacteria or virus infections, which consisted of LB for bacteria and cell culture medium for virus. *Drosophila* larvae were removed at around late second instar/early third instar from the 50mm cornmeal plates, and washed in PBS. Twenty larvae were transferred at a time to a piece of filter paper. The prepared Tungsten needle was dipped into the prepared culture and used to prick the larva. This was repeated for each larva. We then gently brushed water through the line of 20 larvae for easier removal from the filter paper and transferred the larvae onto the fly food in a glass vial.

### RNA extraction, cDNA preparation, and quantitative polymerase chain reaction (qPCR)

Fifteen *Drosophila* larvae were used for RNA extraction for each treatment using the total RNA extraction protocol using the chloroform isopropanol TRIzol method following the manufacturer’s protocol (Invitrogen #15596026). *DiptericinB* and *CecropinA* were targeted with qPCR to confirm the infection with *E.coli*. *DptB* primers: fwd-“TTCTCGAGTGCCTGGGCTTA” and rev-“ATTGGGAGCATATGCCAGTG”, and *CecA*1 primers: fwd-“ TTGGCAAGAAAAATCGAACGC” and rev-“GCTTGTTGAGCGATTCCCAG”. DCV infection was detected using primers specifically targeting a region of the DCV genome: fwd-“GACACTGCCTTTGATTAG” and rev-“CCCTCTGGGAACTAAATG”.

### Quantitative Polymerase Chain Reaction (qPCR)

One μl of RNA per sample was reverse-transcribed with Promega GoScript reverse transcription system (Promega #A5001), using random hexamer primers. The complementary DNA (cDNA) samples were then diluted 10-fold with nuclease free water for qPCR. qPCR was performed on Applied Biosystems StepOnePlus system (ThermoFisher Scientific #4376600) or using Applied Biosystems QuantStudio5 system (ThermoFisher Scientific #A34322), with SensiFast SYBR Hi-ROX mix (Bioline #BIO-92020).

### Localized *lectin-24A* expression microscopy

Larval fat bodies were dissected from 3^rd^ instar *D. melanogaster* larvae. The whole fat body was obtained by first removing the head of the larva then pulling back the cuticle to expose the fat body and other inner tissues. The gut, salivary gland, and other tissues were then removed and discarded. Each isolated fat body is transferred into an Eppendorf tube containing ice-cold PBS. The fat bodies were incubated in 3.7% formaldehyde diluted in PBS for 20 minutes with gentle agitation on an orbital shaker, with the tubes covered in foil. The formaldehyde was then removed by pipetting and washed with ice-cold PBS for three times. Then we again washed the fat bodies two times for 30 minutes each with 0.3% ice-cold PBT with gentle agitation on an orbital shaker, with the tubes covered in foil. The fat bodies were then washed for three times again with ice-cold PBS by pipetting. We then stained the nuclei with 1μg/ml Hoescht staining solution diluted in PBS for 5 minutes, while making sure the staining solution sufficiently cover the whole fat body. The fat bodies were washed again with PBS for three times, then each transferred individually to a clean Eppendorf tube. Immediately cover the fat body tissue with cold Vectashield mounting media (Vector Laboratories # H-1000-10). Pipette tips with the ends cut were used to transfer each fat body with minimal volume of mounting media on to a microscope slide. The fat body was then spread out on the microscope slide using two nickel plated pin holders holding 0.1mm diameter pins. We then slowly added a cover slip to the slide, making sure there were no bubbles. The fat bodies were then imaged using a Leica upright fluorescence microscope (Leica # DM6000M).

### Larval fat body posterior vs. anterior RNAseq dissection

Larval fat bodies were dissected according to the protocol in the previous section “**Localized *lectin-24A* expression microscopy protocol**”. The anterior and posterior of the fat body were separated along the representative line as shown in Figure 2A.

### Library preparation for RNA sequencing

At 24 hpi, groups of 10 male larvae from the stock expressing the *lectin-24A* promoter reporter were dissected, with the anterior and posterior larval fat body sections separated. Age-matched uninfected fat bodies were also dissected and separated into anterior and posterior sections. RNA was isolated from each sample of pooled fat body sections, where for each sample the PBS supernatant was removed and 250μl of Tri-reagent (Ambion #10296010) was added. The fat bodies were homogenized using TissueLyzer II (Qiagen #85300) for 2 minutes at 30Hz. 62.5μl of chloroform was added to each sample and the tubes were shaken for 15 seconds followed by 3 minutes of incubation at room temperature. The samples were then centrifuged for 10 minutes at 12,000xg at 4°C. 66μl of the upper aqueous phase was transferred to a fresh Eppendorf tube and 156μl of propan-2-ol was added. The tubes were then inverted 5 times to mix thoroughly. Following a 10-minute incubation at room temperature, the samples were centrifuged at 12,000xg at 4°C for 10 minutes. The supernatant was removed by pipetting, watching out for small white pellets at the bottom of the tubes, and 250μl of ice-cold 70% ethanol was added. The samples were centrifuged again at 4°C at 12,000xg for 2 minutes and the ethanol was removed. The RNA pellets at the bottom of the tubes were allowed to air-dry very briefly and 20μl of nuclease-free water was added. The RNA pellets were dissolved by incubating the tubes on a heat block adjusted to 45°C for 5 minutes. RNA was quantified using a Qubit RNA HS assay kit (Thermofisher #Q32852). RNAseq libraries were then prepared using a NEBNext Ultra II Directional RNA Library Prep Kit for Illumina (New England Biolabs #E7760S) with NEBNext Multiplex Oligos for Illumina (96 Unique Dual Index Primer Pairs) (New England Biolabs #E64460S) and polyA enrichment module (NEB #E7490S). Up to 1μg of total RNA was used to make each RNAseq library, where mRNA was fragmented at 94°C for 15 minutes following polyA enrichment, according to the protocol recommendations. The adaptor concentrations and number of amplification cycles used were adjusted based on the amount of starting RNA. The quantity in each prepared library sample was measured using Qubit DNA HS assay kit (Thermofisher #Q32851). The library quality was assessed using a Bioanalyzer high sensitivity kit on an Agilent 2100 Bioanalyzer (Agilent #5067-4626). The libraries were sequenced at Source BioScience using Illumina NovaSeq 150bp paired-end reads.

### Statistical analyses of RNAseq data

We used Trim Galore (https://www.bioinformatics.babraham.ac.uk/projects/trim_galore/) for trimming raw RNAseq reads, setting a Phred score of 20 for quality control. Reads with fewer than 50 bases after trimming were removed. The reads were then mapped and counted using STAR v2.6.0 [91] to *D. melanogaster* reference genome Dmel-r6.40. The R package edgeR (RRID:SCR_012802) was used for differential expression analyses. Genes that had a count per million (CPM) above ten in at least four libraries were kept, serving as a threshold for identifying genes with detectable expression in our dataset. Dispersions were estimated using the Cox-Reid profile-adjusted likelihood (CR) method in edgeR, and the expression data were fitted with a negative binomial generalized linear model (GLM). Differential expression of genes was then determined with a quasi-likelihood (QL) *F*-test, where a false discovery rate (FDR) of 0.05 was set as the significance threshold.

### Gene Ontology enrichment analysis

A Gene Ontology (GO) enrichment analysis was conducted using the R package goseq [92]. The list of genes detected in RNAseq analysis was used as a background set. Sets of significantly upregulated and significantly downregulated genes with an FDR < 0.05 and up or down-regulation of at least one logFC were then analyzed separately to determine GO enrichment. The length bias inherent to RNAseq data were accounted for by calculating a Probability Weighting Function (PWF) using goseq, that gives a probability that a gene will be differentially expressed based on its length alone. A null distribution for GO category membership was approximated with Wallenius distribution, and each GO category is then tested for over and under representation amongst the set of differentially expressed genes. Where there are more than 5 GO terms in any of the three major Gene Ontology branches (Biological Processes, Cellular Components, and Molecular Functions), the list of GO terms was summarized using REVIGO [93] (RRID:SCR_005825).

### Yeast-1-hybrid (Y1H)

The Y1H analysis of the *lectin-24A* upstream region was carried out following the system developed in [94].

### RNAi knockdown assay

We used fly stocks from the Transgenic RNAi Project (TRiP), which consisted of flies that utilize the GAL4/UAS driven system to silence a specific gene using short hairpin RNA and RNA interference (RNAi). To perform the experiment, a fly stock that expresses a *daughterless* driven GAL4 (*da*-Gal4), a temperature-sensitive tubulin driven GAL80 (*tub*-Gal80^ts^), and the *lectin-24A* promoter Venus reporter (LP437) was crossed to the RNAi-expressing line targeting *Stat92E* (BDSC#33637). The cross generated F_1_ progeny with the genotype LP437/+; tub-Gal80ts/+; da-Gal4/UAS-TRiP. Egg lay was carried out at 25°C overnight, then the developing larvae were reared at 29°C. On the 4th day, larvae were infected for 3 hours with 3 female wasps in each vial, and larvae were collected 24 hpi for protein extraction to analyze fluorescence expression of *Venus^LP437^*, as stated in “**Lectin promoter reporter assay**”.

### Truncated *lectin-24A* promoter fluorescence reporter

We created three truncated versions of the *lectin-24A* promoter fluorescence reporter, starting 20bp 5’ of each of the three indels in the *lectin-24A* upstream region. The constructs were made in the same way as the original *lectin-24A* fluorescence reporter construct [12]. We digested pWallium20 backbone plasmid with the restriction enzymes HpaI and PstI (NEB # R0105S and R0140T) followed by gel purification. The forward primers used to amplify each promoter sequence are listed in the following table. The reverse primer used to amplify each sequence was 5’- ATTATAAGCTGCAATAAACAAGTTTTACTTGTACAGCTCGTCCATGCC-3’. The insert sequence is amplified from the full-length *lectin-24A* promoter fluorescence reporter, and it includes both the truncated promoter sequence as well as the Venus open reading frame. The digested pWallium20 backbone and the amplified PCR fragment were assembled using a HiFi assembly reaction (NEB #E5520), with an incubation time of 1 hour at 50 °C. The transformation plasmids were then purified by miniprep (Qiagen # 27104). Plasmid sequences were then verified by Sanger sequencing using the primers M13 Reverse 5’- CAGGAAACAGCTATGAC-3’ and the reverse primer to amplify the insert sequence. All constructs were injected into *D.melanogaster* line attP2Ar5, harboring an attP landing site on the X chromosome (BDSC#24480) by Bestgene Inc, USA, and transformants were selected by eye color.

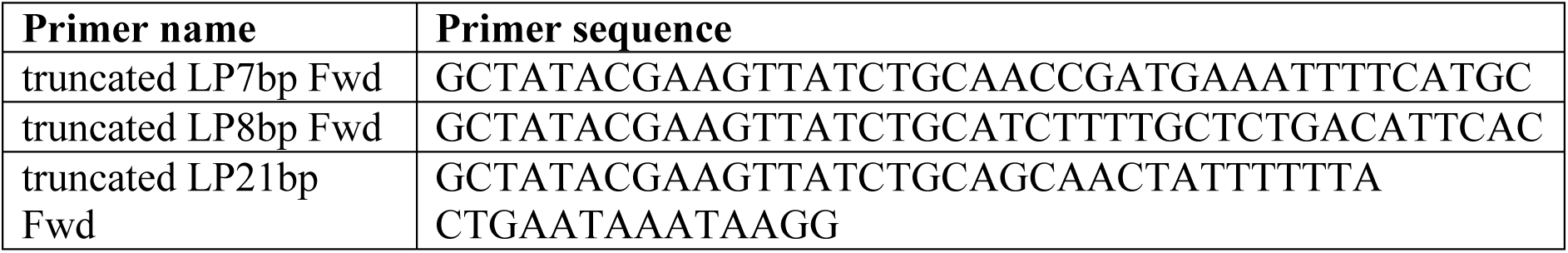

### Scrambling TFBMs with site-directed mutagenesis (SDM)

To generate *lectin-24A* promoter fluorescence reporter with scrambled transcription factor binding sites, we used Q5 site-directed mutagenesis kit (NEB# E0554S). Primers were designed according to the sites to be scrambled as shown in the table below, where the lowercase letters represented changes made to the original promoter sequence. The truncated LP8 plasmid was used for scrambling instead of the full-length *lectin-24A* fluorescence promoter reporter.

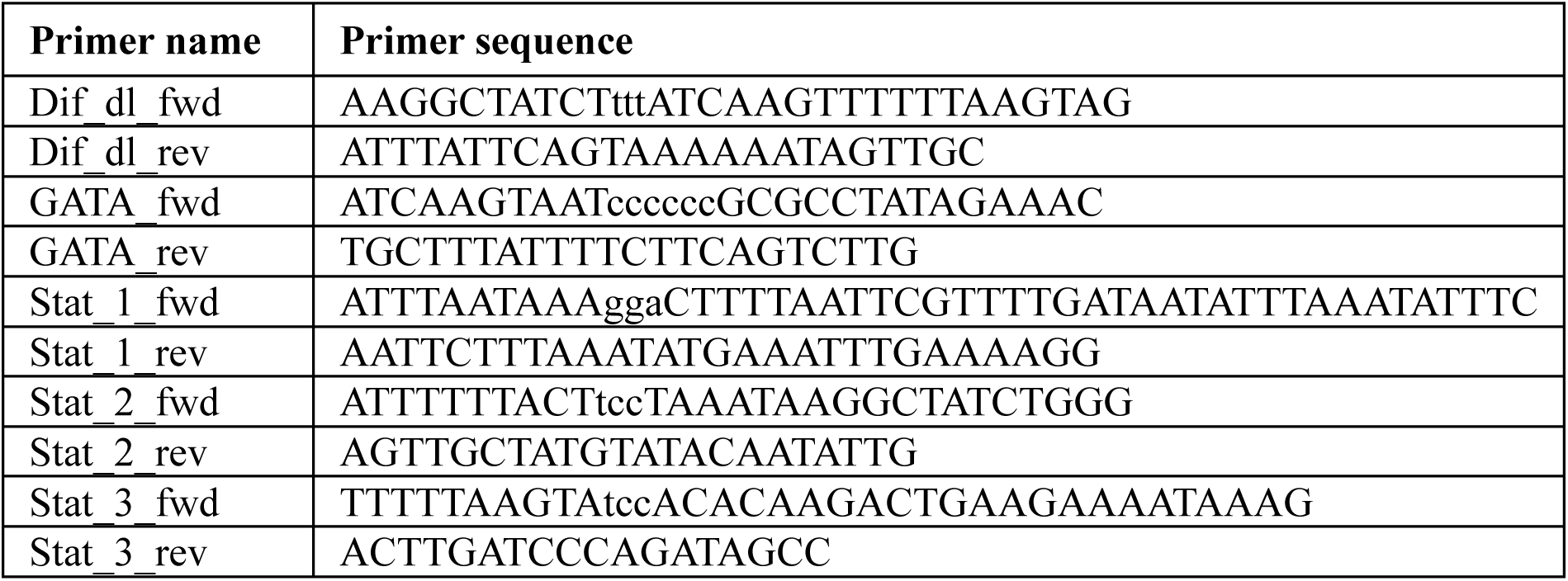

### JAK/STAT pathway *et* mutant and *lectin-24A* reporter recombination

The *et* mutant line is a kind gift from Michèle Crozatier [21]. The *lectin-24A* promoter fluorescence reporter was recombined with the *et* mutant, as both the construct and the gene are on the X chromosome. A line with attached-X chromosomes (genotype: C(1)DX, y w f/y cv v shi[ts1] f mal/y[+]Y) from the Genetics Department at the University of Cambridge was used to follow chromosomes with recombination events. Females from the *lectin-24A* promoter fluorescence reporter line was first crossed with *et* mutant line males, then the female offspring were crossed to *lectin-24A* promoter fluorescence reporter males. Individual male offspring of this cross, which might carry a recombined X chromosome are crossed to females of the attached-X line. As the X chromosomes of males are always inherited through the male offspring when crossed to an attached-X female, any recombined X chromosomes in the males are kept in males.

Male offspring from each of the cross above were first screened for fluorescence using an epifluorescence microscope. Stocks that show Venus fluorescence were then screened for *et* mutation by PCR. Genomic DNA was extracted from males of each stock. Two pairs of primers from [21] were used to check *et* mutation. Fwd-1 & Rev-1 give an amplicon of 1186bp, and Fwd-2 & Rev-2 give an amplicon of 526bp. Stocks carrying the *et* mutation would not give a PCR product from either of these primer pairs. Male larvae from stocks that have successfully recombined were used in subsequent assays for *lectin-24A* promoter fluorescence.

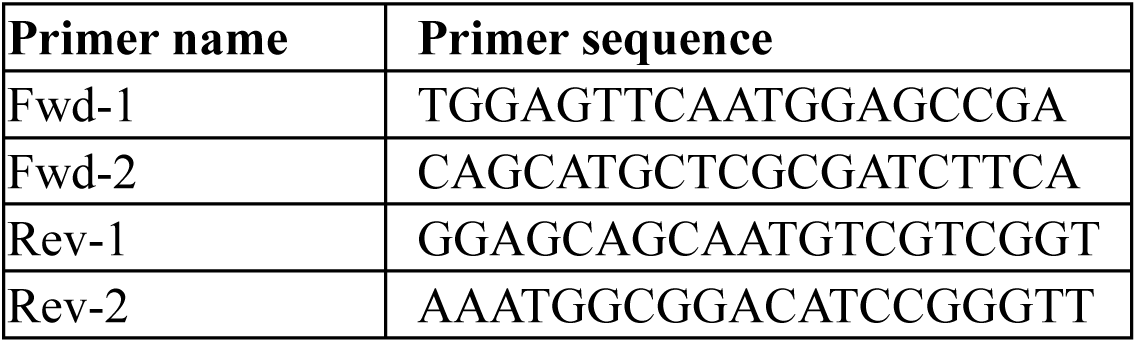

### DomeΔCYT assay

The stock expressing *DomeΔCYT* under UAS control is a kind gift from Michèle Crozatier. We crossed females expressing the larval fat body driver *Cg-GAL4* or the ubiquitous *da-GAL4* driver, as well as *tub-GAL80^ts^*, with males expressing *UAS*-*DomeΔCYT*. The stock also carried *UAS-et* with a balancer chromosome that could not be identified in larvae, so a portion of the F_1_ progeny also expressed *UAS-et*. The control cross was with females expressing only the *Venus^LP437^* reporter with no driver. We carried out egg lay and allowed the larvae to develop at 18°C, until the 7^th^ day after egg lay, when we moved the larvae to 29°C 3 hours prior to infection with parasitoids. Sample collection was performed at 24 hpi. The assays were then carried out as stated in “**Lectin promoter reporter assay**”.

### *hop^Tum^* assay

Females expressing the *Venus^LP437^* reporter were crossed with males of the *hop^Tum^* genotype. Larvae were allowed to develop at 25°C. Since both the reporter and *hop* are located on the X chromosome, only female F_1_ heterozygotes were selected during sample collection at 24 hpi. The assays were then carried out as stated in “**Lectin promoter reporter assay**”.

### Dominant positive *Pnr* mutant expression in the *Drosophila* fat body

We expressed the dominant positive mutant *Pnr^D4^* under the control of UAS (BDSC#36546) in the *Drosophila* larval fat body using the driver *Cg-GAL4*. The larvae were allowed to develop at 25°C until 3 hours prior to infection with parasitoids, then they were moved to 29°C until sample collection at 24 hpi, as stated in “**Lectin promoter reporter assay**”.

### Somatic mutagenesis of key immune pathway genes

In order to create somatic knockouts of immune pathway genes, we created a line that expresses the *lectin-24A* promoter reporter and *Act5c-Cas9*. We crossed virgins of this line to males expressing guide RNAs (gRNAs) targeting each gene. The gRNA-expressing lines used are listed below. Primers used for PCR and sequencing to confirm the somatic mutagenesis are also listed. The assays were carried out as stated in “**Lectin promoter reporter assay**”.

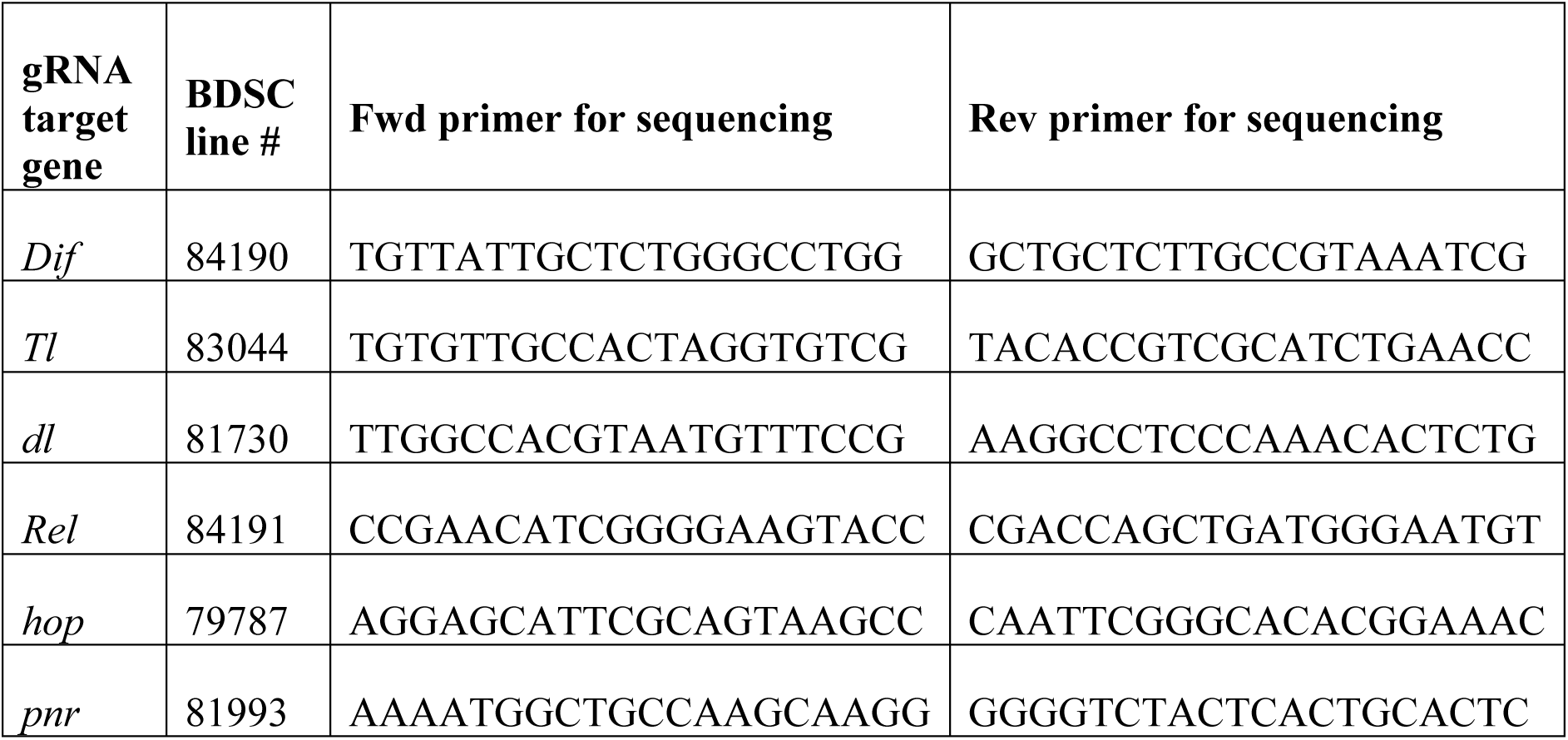

### Overexpression of NF-κB transcription factors dl and Rel

The stocks for expressing *dl* (FlyORF Stock# F000638) and *Rel* (FlyORF Stock# F001780) under UAS control were obtained from the Zurich ORFeome Project [95]. We crossed females expressing the larval fat body driver *Cg-GAL4* and *tub-GAL80^ts^*, with males expressing *UAS*-*dl* or *UAS*-*Rel*. The control cross was with females expressing *Venus^LP437^* with no GAL4 driver. Egg lay and larval development were carried out at 25°C until 3 hours prior to wasp infection on the 4^th^ day after egg lay, when they were moved to 29°C, and sample collection was done at 24 hpi, following “**Lectin promoter reporter assay**”. The UAS-dl stock carried a balancer that could not be identified at the sample collection stage in larvae, so the larvae collected from the UAS-dl crosses were with a mix of *Cg-GAL4/UAS-dl* and *Cg-GAL4/+* genotypes.

### Enrichment of TFBMs in the upstream region of parasitism-induced genes

FIMO from the MEME suite version 5.5.5 was used to scan sequences for occurrences of TFBMs [96,97]. Default parameters were used for GATA TFBMs, and a significance threshold of 0.005 was used for NF-κB and STAT92E TFBMs. MEME format position weight matrices (PWMs) were generated from binding sites collected in flyfactorsurvey [98] for GATA sites, and from IUPAC motifs for NF-κB and STAT sites (Figure 4C).

## Data availability

The RNAseq data has been submitted to the NCBI Sequence Read Archive under the BioProject number PRJNA1021619, with BioSample accessions from SAMN37565424 to SAMN37565439. Processed data files and scripts to analyze data are available on Apollo, the University of Cambridge repository (https://doi.org/10.17863/CAM.108488).

