## Supplementary file for "A humoral immune response to parasitoid wasps in *Drosophila* is regulated by JAK/STAT, NF-κB and GATA"

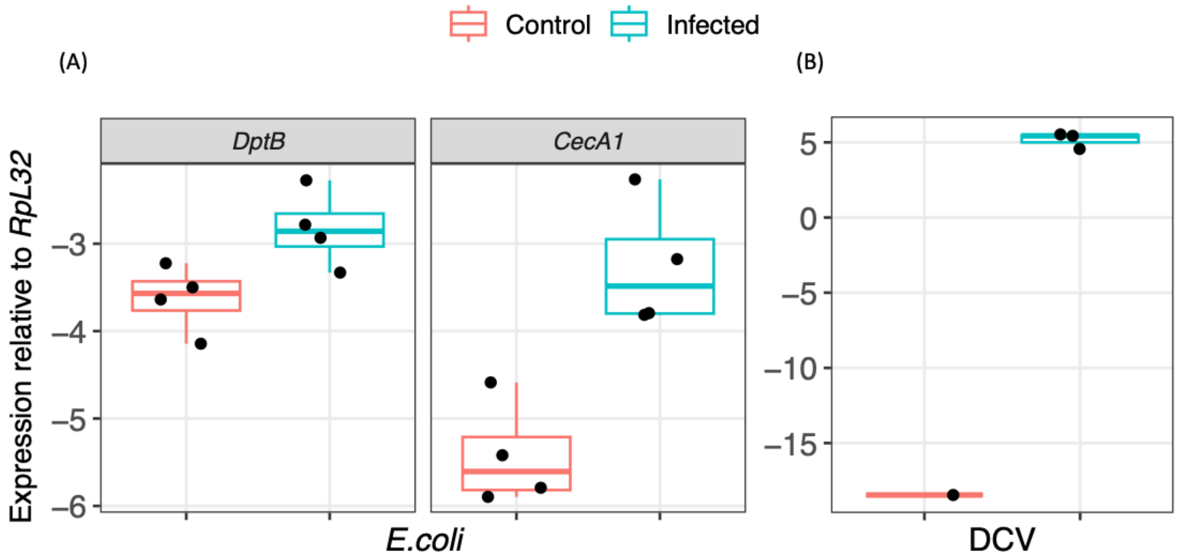


**Supplementary Figure 1. Bacteria and virus infection of *Drosophila* larvae.** (A) Induction of antimicrobial peptides, *DptB* and *CecA1*, following infection of larvae with *E.coli* by pricking at 6 hours post-infection (hpi). (B) Detection of *Drosophila* C Virus (DCV) in larvae infected with DCV at 24 hpi, using primers amplifying a region of the DCV genome. Each point represent a pool of 15 larvae.

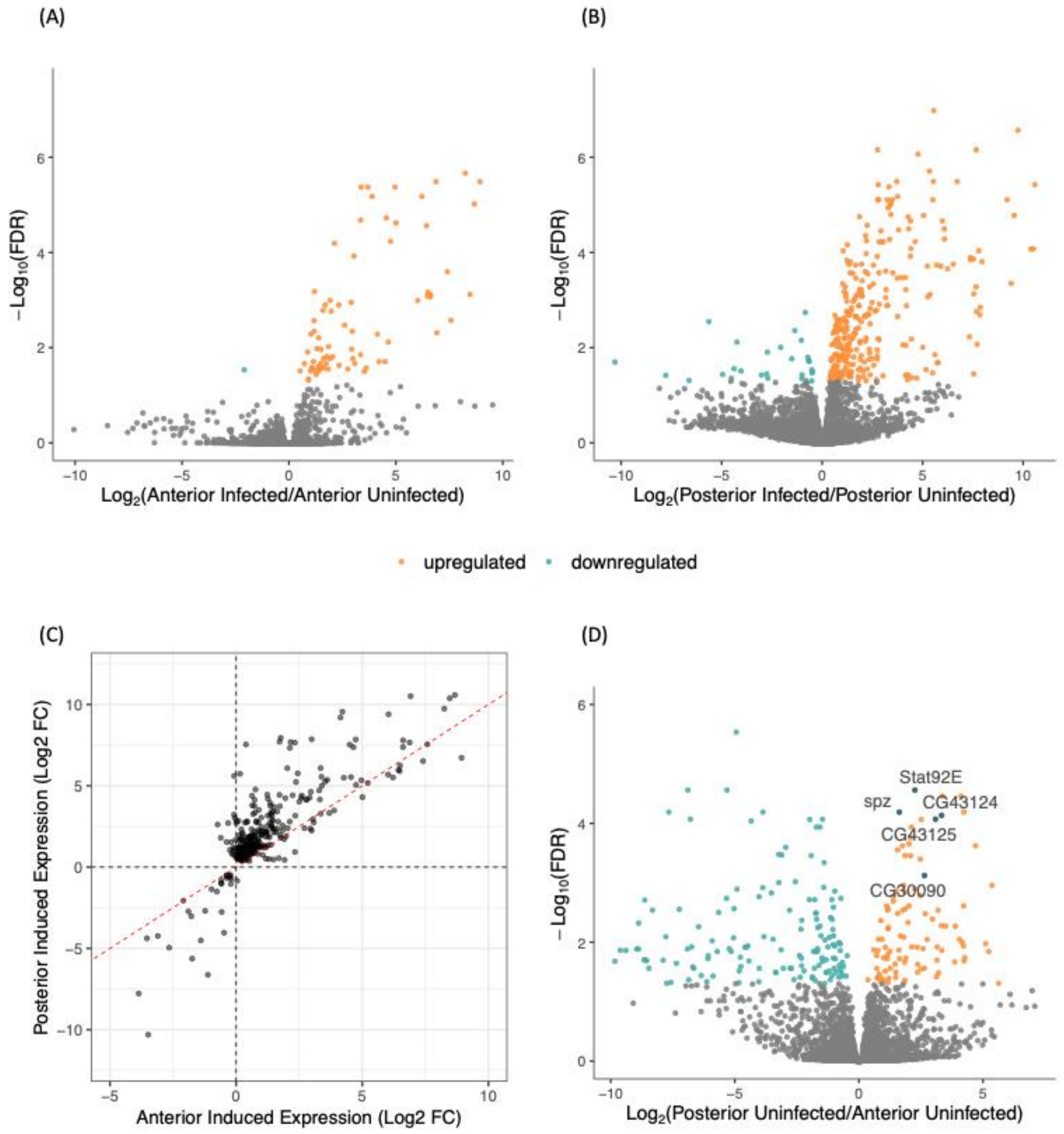


**Supplementary Figure 2. RNA sequencing of *Drosophila* larval fat body sections.** (A) Differential expression in the anterior section of the larval fat body in response to parasitization at 24 hours post-infection (hpi). (B) Differential expression in the posterior section of the larval fat body in response to parasitization at 24 hpi. (C) Scatterplot showing gene expression changes ($log_2FC$) in the posterior vs. anterior of the fat body at 24 hpi. (D) Differential expression between the anterior and posterior larval fat body sections under uninfected condition.

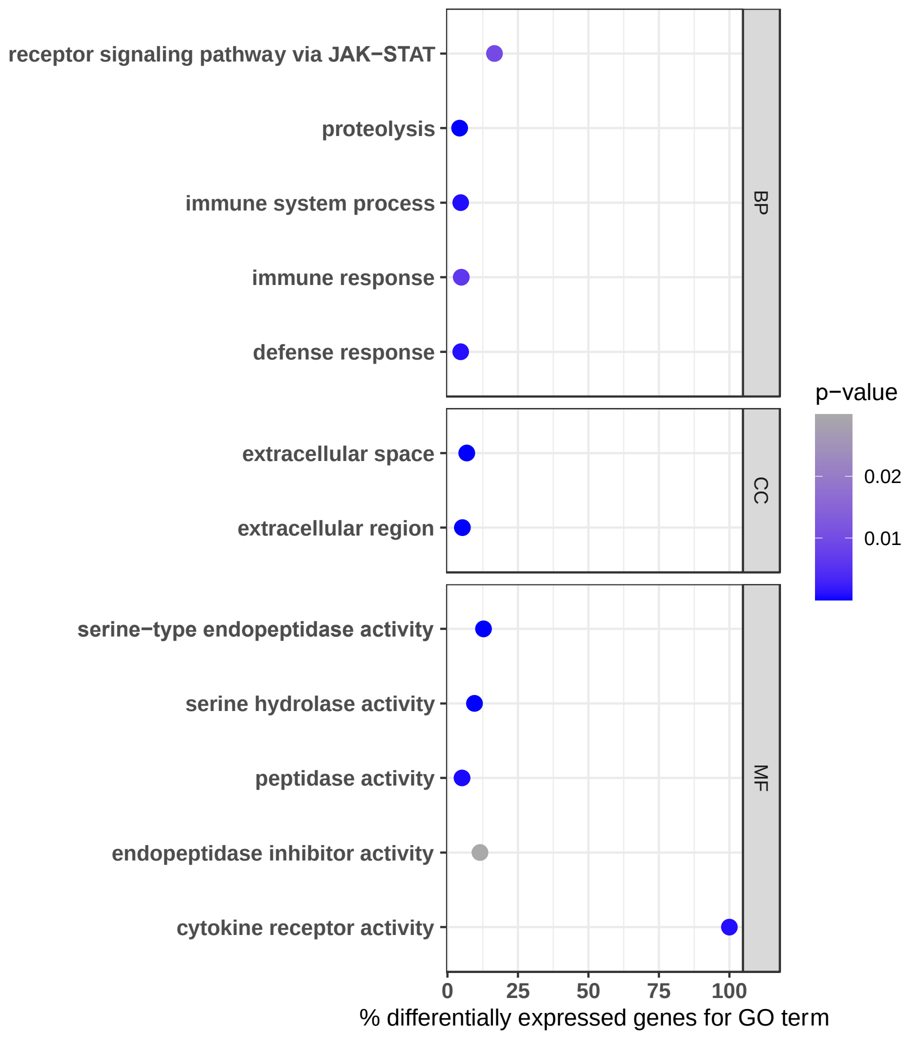


**Supplementary Figure 3. Gene ontology (GO) enrichment for significantly upregulated genes in the posterior of the fat body compared to the anterior at 24 hours-post infection.** BP – Biological Process, CC – Cellular Components, MF – Molecular Functions.

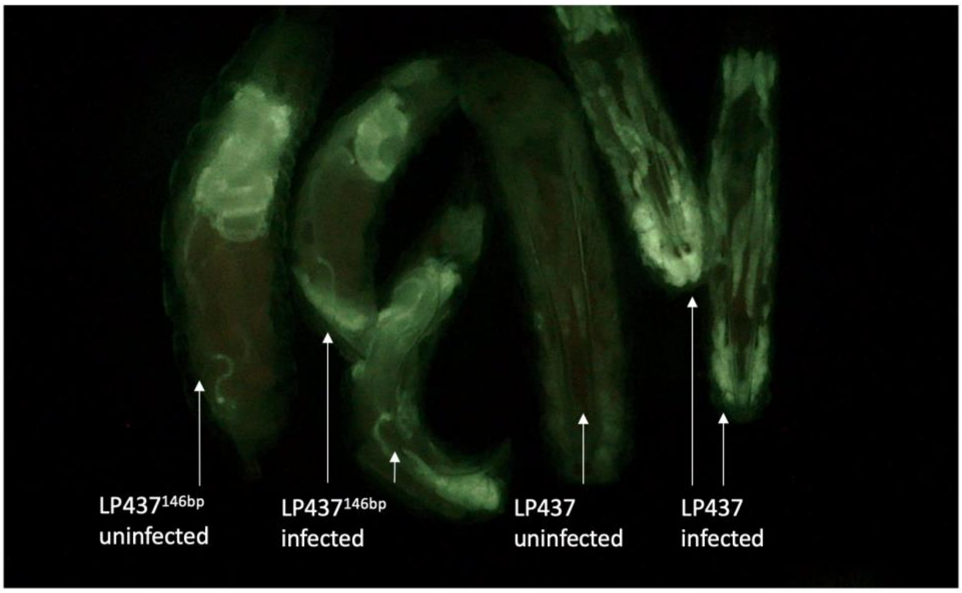


**Supplementary Figure 4. Truncated Venus reporter construct expression in larvae.** Expression of Venus driven by the truncated (LP437$^{146bp}$) or full-length upstream sequence of *lectin-24A*, imaged at 24 hpi under GFP filter.

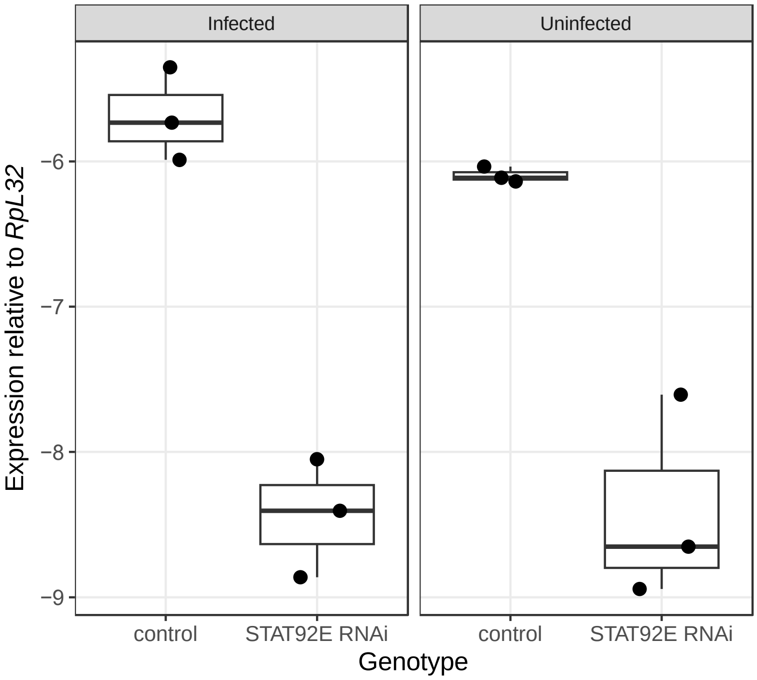


**Supplementary Figure 5. *Stat92E* RNAi knockdown efficiency.** Level of *Stat92E* expression between control and *Stat92E* knockdown larvae. F1 larvae assayed were produced from a cross between females expressing a ubiquitous *da-GAL4* driver and males expressing double-stranded RNA for RNAi of *Stat92E* under UAS control. The control is using males of the same genetic background but expressing *UAS-GFP*.

(A)

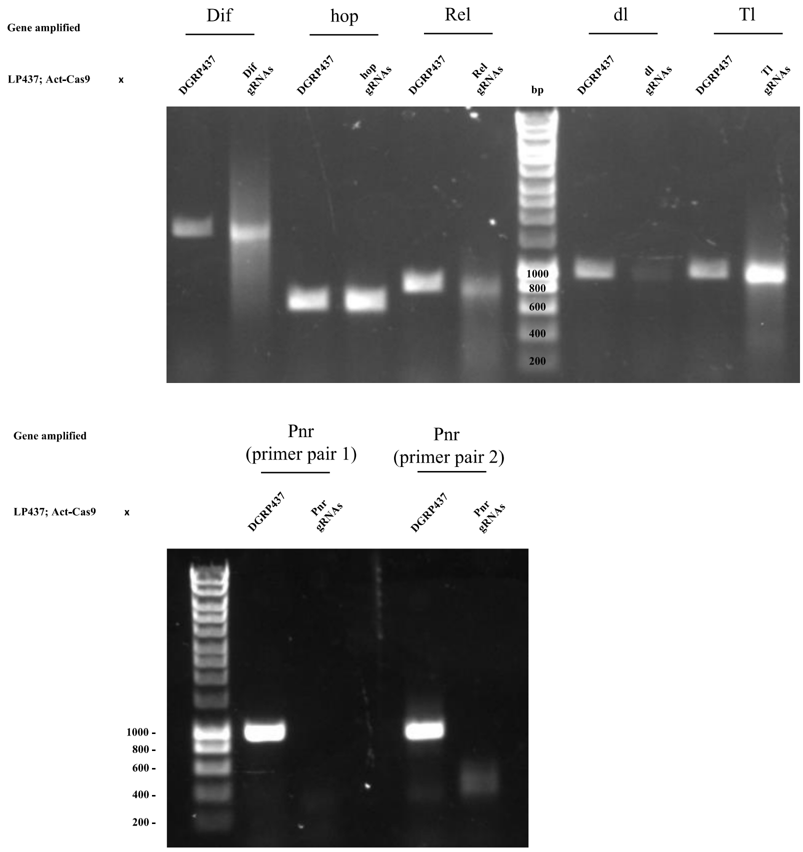
Gene amplified
Dif
hop
Rel
dl
Tl
LP437; Act-Cas9 x
DGRP437
Dif gRNAs
DGRP437
hop gRNAs
DGRP437
Rel gRNAs
bp
DGRP437
dl gRNAs
DGRP437
Tl gRNAs
1000
800
600
400
200
Gene amplified
Pnr (primer pair 1)
Pnr (primer pair 2)
LP437; Act-Cas9 x
DGRP437
Pnr gRNAs
DGRP437
Pnr gRNAs
1000 -
800 -
600 -
400 -
200 -

(B)

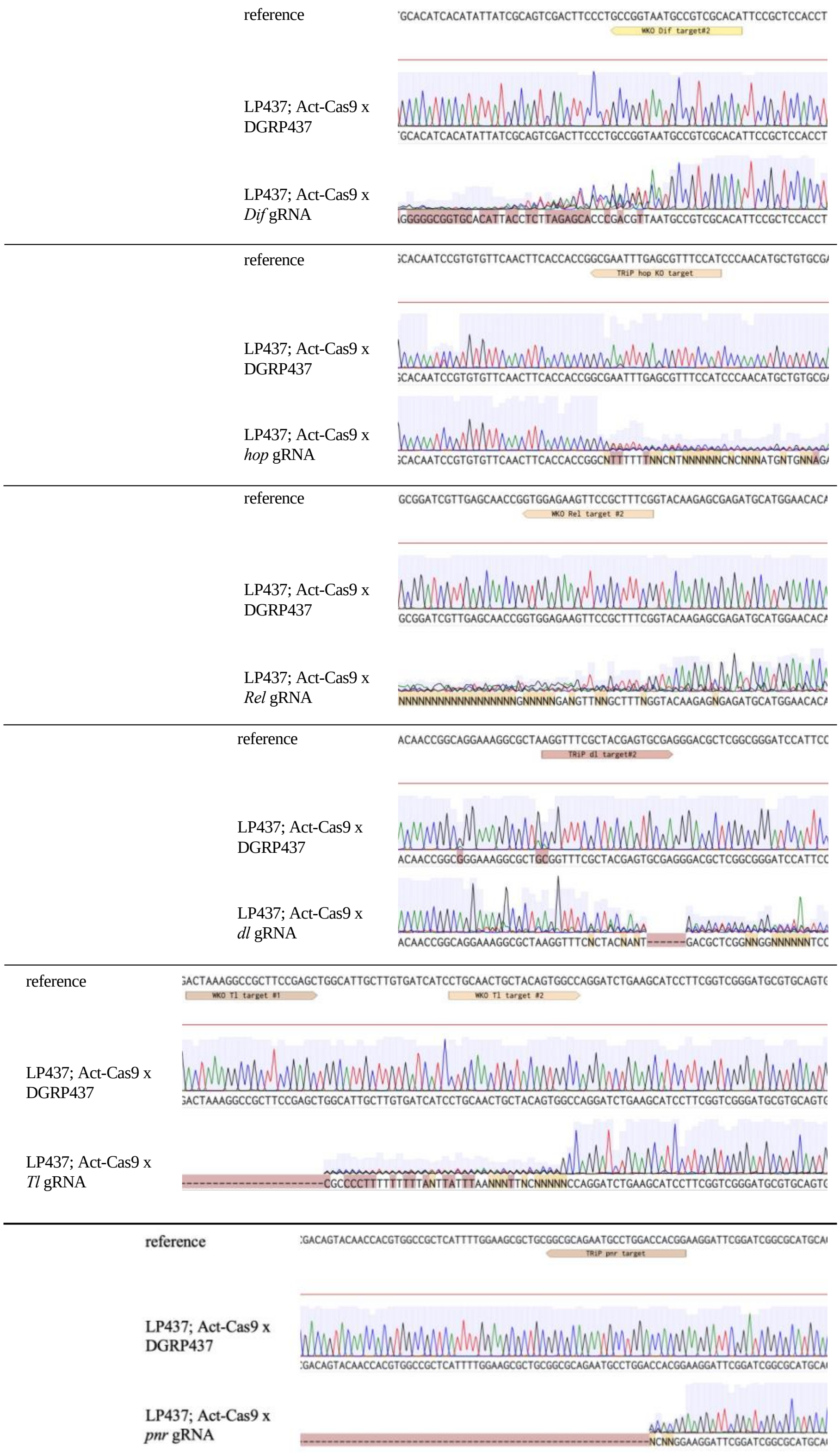


**Supplementary Figure 6. Somatic mutagenesis of immune pathway genes in F$_1$ heterozygotes ubiquitously expressing guide RNAs targeting each gene and Cas9 protein.** (A) Gel of PCR products from amplifying the open-reading frame using pooled genomic DNA of 5 adult flies from the F$_1$ progeny of lines expressing gRNAs targeting each gene crossed to a line ubiquitously expressing Cas9, compared to control larvae. (B) Alignment of Sanger sequencing results of the PCR products amplified from control larvae and larvae with somatic mutagenesis, compared to the reference sequence.

| Transcription factor | Motif sequence |
|---|---|
| pnr | HGATAASV |
| srp | WGATAASV |
| GATAd | TGATAASV |
| GATAe | HGATAAS |

**Supplementary Table 1. Binding motif sequences of GATA pathway transcription factors.**

| STAT92E TFBM | STAT92E predicted binding site upstream of *lectin-24A* | Position relative to TSS | Sequence post-scramble |
|---|---|---|---|
| TTC(N)$_5$AA | TTCCTTTTAA | -200 | **GGA**CTTTTAA |
| | TTTTACTGAA | -137 | TTTTACT**TCC** |
| | TTAAGTAGAA | -99 | TTAAGTA**TCC** |
| TTC(N)$_4$AA | TTTAAAGAA | -221 | - |
| | TTTACTGAA | -136 | TTTACT**TCC** |
| TT(N)$_5$AA | TTTAATAAA | -209 | - |
| | TTTTGATAA | -186 | - |

**Supplementary Table 2. Predicted STAT92E binding sites in the *lectin-24A* upstream region with the relative position of each site and the sequence after scrambling. The scrambled positions are indicated with bold and underline.**
